# Minor changes in electrostatics robustly increase VP40 membrane binding, assembly, and budding of Ebola virus matrix protein derived virus-like particles

**DOI:** 10.1101/2024.01.30.578092

**Authors:** Balindile B. Motsa, Tej Sharma, Prem P. Chapagain, Robert V. Stahelin

## Abstract

Ebola virus (EBOV) is a filamentous negative-sense RNA virus which causes severe hemorrhagic fever. There are limited vaccines or therapeutics for prevention and treatment of EBOV, so it is important to get a detailed understanding of the virus lifecycle to illuminate new drug targets. EBOV encodes for the matrix protein, VP40, which regulates assembly and budding of new virions from the inner leaflet of the host cell plasma membrane (PM). In this work we determine the effects of VP40 mutations altering electrostatics on PM interactions and subsequent budding. VP40 mutations that modify surface electrostatics affect viral assembly and budding by altering VP40 membrane binding capabilities. Mutations that increase VP40 net positive charge by one (e.g., Gly to Arg or Asp to Ala) increase VP40 affinity for phosphatidylserine (PS) and PI(4,5)P_2_ in the host cell PM. This increased affinity enhances PM association and budding efficiency leading to more effective formation of virus-like particles (VLPs). In contrast, mutations that decrease net positive charge by one (e.g., Gly to Asp) lead to a decrease in assembly and budding because of decreased interactions with the anionic PM. Taken together our results highlight the sensitivity of slight electrostatic changes on the VP40 surface for assembly and budding. Understanding the effects of single amino acid substitutions on viral budding and assembly will be useful for explaining changes in the infectivity and virulence of different EBOV strains, VP40 variants that occur in nature, and for long-term drug discovery endeavors aimed at EBOV assembly and budding.

## Introduction

The Ebola virus (EBOV) is a filamentous negative sense RNA virus which causes severe hemorrhagic fever. EBOV is one of the most dangerous known pathogens with a high fatality rate. Multiple outbreaks of EBOV have occurred since the 1970s with the most widespread outbreak starting in December 2013. This outbreak continued through May of 2016 and had a fatality rate of approximately 50%. Smaller EBOV outbreaks are recurrent especially in regions of West Africa because the virus is present in animal reservoirs. As of March 2023, there have been ∼35,000 total cases of EBOV and 15,000+ deaths with fatality rates ranging from 30%-90% (1). Currently, there is only one FDA approved vaccine for the prevention of EBOV and two monoclonal antibody treatments (2) that are effective against the EBOV glycoprotein (GP) and prevent viral entry (3, 4). Despite this success, these therapeutics may only be effective against the Ebola Zaire strain, one of six known strains of EBOV. Further, there are still no FDA approved small molecule treatments for those infected with EBOV. Despite multiple EBOV outbreaks in the last decade, we still lack a clear understanding of how new viral particles are formed in infected cells and spread through virus assembly, budding, and release. Deeply understanding the EBOV assembly and budding processes will illuminate drug targets in the virus life cycle (5).

The EBOV genome encodes for seven proteins, each with distinct functions in the viral lifecycle (6). Of these proteins, the EBOV matrix protein VP40 has emerged as an attractive target for therapeutic intervention (5, 7). VP40 is one of the most conserved viral proteins and inside the cell VP40 rearranges into different structures, each with a distinct function that is required in the viral lifecycle (8, 9). VP40 forms an RNA binding ring octamer that is necessary for the transcriptional regulation of viral gene expression during an infection (8, 9). VP40 forms a dimer that is necessary for the assembly and budding of new virions from the inner leaflet of the host plasma membrane (PM) (8, 10, 11). When transiently expressed in cells, the VP40 dimer is trafficked to the inner leaflet of the PM where it forms higher order oligomeric structures via interactions between the dimers (11). This PM localization and VP40 filament formation in-turn allows for the budding and release of filamentous virus-like particles (VLPs) (12–14). These VLPs are not infectious, however, they are similar in size and nearly indistinguishable from the authentic virus (13, 15, 16). Co-expression of VP40 with the EBOV GP produces VLPs that can be used to model viral entry in a BSL-2 lab setting (17, 18).

VP40 harbors an N-terminal domain (NTD) and a C-terminal domain (CTD) (19, 20). NTD interactions (predominantly through an alpha-helical interface) are responsible for the formation and stability of the VP40 dimer (8, 21) in the cytoplasm, and CTD interactions are important for trafficking of the dimer (22, 23) and binding anionic PM lipids (19, 20, 24). VP40 binds to anionic lipids in the PM inner leaflet via electrostatic interactions to hijack the host membrane and form the viral lipid envelope (10, 12, 14, 20, 24, 25). VP40 lipid interactions are critical for PM association and VP40 oligomerization (10, 12) to form a matrix layer along the membrane interface. The VP40 dimer traffics to the inner leaflet of the PM (8) where it interacts with the PM through key lysine residues (especially Lys^221^, Lys^224^, Lys^225^, Lys^274^, and Lys^275^) in its C-terminal domain (CTD) (8, 26). Once there, electrostatic interactions trigger the rearrangement of VP40 dimers into hexamers and larger oligomers (27–29), which construct a filamentous matrix structure that is critical for budding (11). The binding of VP40 to the PM is mediated by the presence of phosphatidylserine (PS) and phosphatidylinositol 4,5-bisphosphate (PI(4,5)P_2_) in the membrane. VP40 has been shown to bind PS containing vesicles (30, 31) and this binding is important for PM association and the formation on VLPs (10). Additionally, VP40 lipid interactions are sufficient to induce structural changes *in vitro* to lipid membranes containing phosphatidylserine (32). PS promotes the initial binding of the VP40 dimer to the PM and triggers oligomerization (e.g., hexamer formation) (10, 28, 30). PI(4,5)P_2_ is needed for the extensive oligomerization from PS induced hexamers and oligomer stability (28). When lipid binding is abolished, VP40 PM assembly and budding is significantly reduced.

The assembly and trafficking of VP40 to the PM requires a network of protein-protein and lipid-protein interactions (PPIs and LPIs). Studying these interfaces is important for understanding how VP40 structure and function regulates trafficking and assembly of the protein and may identify key sites for target therapeutics to inhibit VP40 function. Previous studies have detailed mechanisms by which VP40 interacts with PS and PI(4,5)P_2_ (10, 26, 33) and that VP40 interactions with these lipids are driven by cationic residues on the surface of the CTD. Further, mutational analysis identified mutations that can more favorably interact with PS via restraint of a loop region (198-GSNG-201) near the lipid binding interface (34). Therefore, we examined in more detail the mechanisms by which cationic charge near the VP40 CTD membrane binding interface influences PM assembly, lipid binding, and budding of VP40.

## Results

### VP40 plasma membrane localization is dependent on C-terminal domain surface electrostatics

The VP40 CTD Lys residues (Lys^224^, Lys^225^, Lys^274^, and Lys^275^) have been shown to mediate interactions between VP40 and PS or PI(4,5)P_2_ containing membranes (33). Electrostatics play a critical role in VP40 membrane association (8, 10, 12, 26, 28, 33), and the overall EBOV budding process (8, 26, 33). To assess the role of surface cationic charge in the VP40 CTD, we first assessed a loop region (198-GSNG-201) previously shown to open and close intermittently in the absence of membrane interactions (34). This loop region is adjacent to key PS and PI(4,5)P_2_-binding residues Lys^224^ and Lys^225^ (26, 33) and is restrained from extending towards the membrane due to interactions between the 198-GSNG-201 loop and Cys^311^ in the CTD (Fig. 1). We hypothesized that increasing cationic charge in this region of the VP40 CTD would increase interactions with the anionic inner leaflet of the PM.

**Figure 1.**
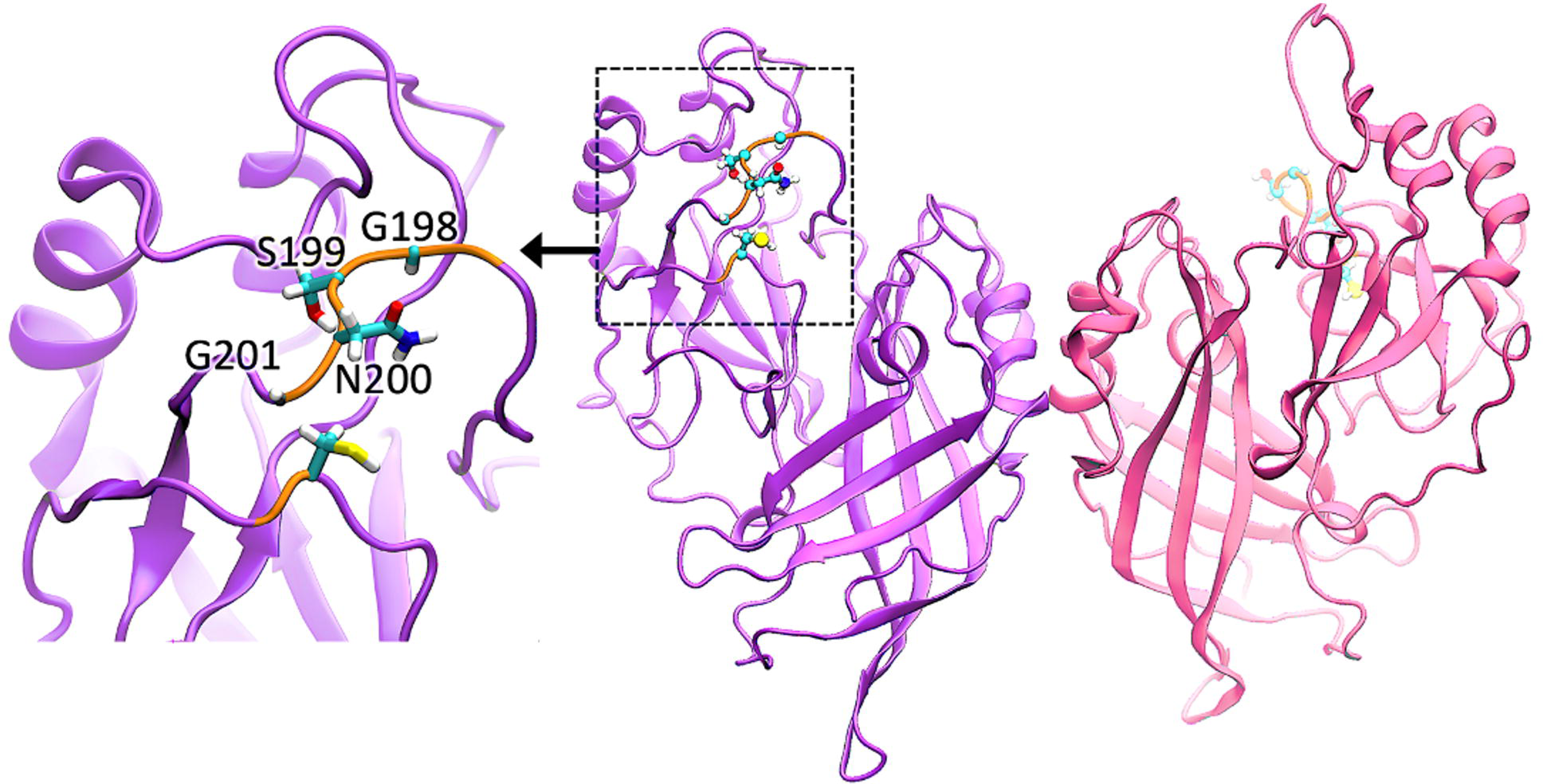
The VP40 dimer structure is shown with the C-terminal domain loop region assessed for electrostatic changes. VP40 (PDB ID: 4ldb) dimer is shown with each protomer in violet and pink, respectively. The protomer on the left highlights the C-terminal domain loop (198-GSNG-201) with stick residues and a zoomed-in view is shown to the left with residues in the loop labeled.

To test this hypothesis, we prepared several mutations in this loop region: G198R, G198A, G198D, and G201R. Localization of WT VP40 and mutations in cells was assessed using a well-established EGFP-VP40 PM localization assay using confocal microscopy (26, 29). WT VP40 localized predominantly to the PM 24 hours post-transfection and formed extensions indicative of pre-assembled VLPs (10). The G198A, G198R and G201R mutations had a similar phenotype to WT localization (Fig. 2) and all three of these mutations exhibited PM localization and formed VLPs that could be seen pre-assembled at the PM (Fig. 2*A*). In contrast, the G198D and L117A mutations displayed no distinct cellular localization but were diffusely distributed between the cytoplasm and the nucleus, like EGFP (Fig. 2A). Quantification of images using ImageJ revealed that the G198A mutant had similar PM localization compared to WT (Fig. 2B). Both the G198R and G201R mutations significantly increased VP40 PM localization while the G198D mutant had decreased PM localization (Fig. 2*B*). The L117A dimerization mutant completely abolished PM localization and exhibited localization like EGFP (Fig. 2*B*). Notably, similar results were obtained after just eight hours of WT VP40 or mutant expression (Fig. S1) with both G198R and G201R exhibiting a significant increase in PM localization. These data suggest mutations that increase overall CTD charge (G198R and G201R) increase VP40 PM association while mutations that decrease (G198D) this charge decrease PM association. Mutations that do not affect the overall charge of VP40 (G198A) do not influence VP40 PM association.

**Figure 2.**
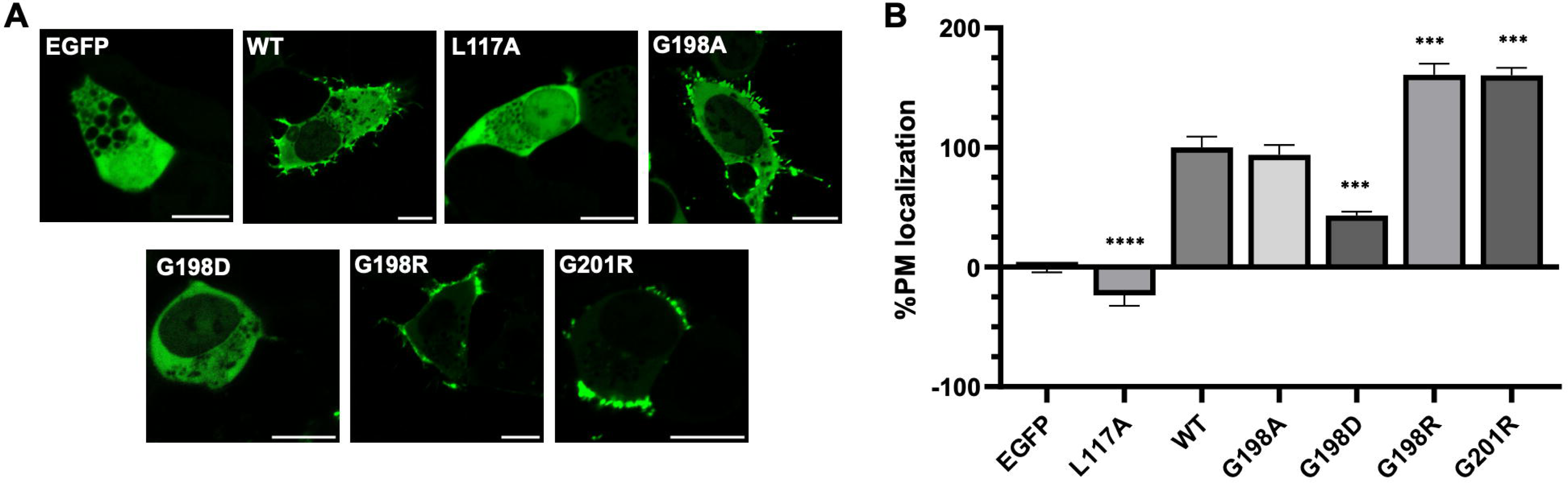
Cellular localization of WT VP40 and respective mutants in HEK293 cells. *A*, Representative images of EGFP, EGFP-WT-VP40, EGFP-VP40-L117A, EGFP-VP40-G198A, EGFP-VP40-G198D, EGFP-VP40-G198R, EGFP-VP40-G201R constructs imaged 24 hours post-transfection. Scale bar = 10 μm. *B*, Average PM localization for each construct. Image analysis was done in Image J to determine percent PM localization (%PM localization) for each construct. %PM localization was defined as (PM intensity/(PM intensity+ cytosol intensity))*100. Data was normalized to %PM localization of WT as 100% localization and %PM localization of EGFP as 0% localization. A WGA-Alex647 PM marker was used to mark the PM in each cell. Error bars are the SEM from three independent experiments with at least 9 images analyzed per replicate. A one-way ANOVA was performed with multiple comparisons compared with WT PM localization (***p<0.001 and ****p<0.0001).

### Increasing cationic charge of the 198-201 loop region enhances budding of virus-like particles

Since changes in electrostatics at positions 198 and 201 affected VP40 PM assembly, we sought to determine the effect of these mutations on VLP production. VLPs produced from each construct were collected 48 hrs post-transfection as described in the methods section. The isolated VLPs and cell lysates were subjected to Western blot for the EGFP tag while using blotting for GAPDH in cell lysates as a loading control. In agreement with the lack of PM localization, the L117A mutant did not produce any detectable VLPs (Fig. 3*A* and *B*). This is also supported by previous studies demonstrating abrogation of the VP40 dimer via a L117R mutation inhibited VLP formation (8). Next, mutation of Gly^198^ to Ala did not significantly change VLP production (Fig. 3*A* and *B*) akin to no changes in PM localization of G198A discussed above. In contrast, introduction of Arg at position 198 or 201 (G198R and G201R), respectively, lead to a significant increase in VLP formation (Fig. 3*A* and *B*) compared to WT VP40. However, introduction of an Asp at position 198 (G198D) lead to a significant reduction in VLP formation (Fig. 3*A* and *B*). Despite the G198D mutation having a low level of PM localization, VLP formation was still possible and within the detection limits of the assay. This contrasts with L117A, where disruption of dimer formation inhibits VLP formation. This suggests that G198D can still form dimers that can undergo CTD-CTD interactions, a prerequisite of effective VLP formation . These results indicate that in addition to altering PM assembly, mutations that alter VP40 overall charge density by ∼ +1 or -1 also greatly affect VLP budding efficiency. Herein an increase in just one positive charge leads up to a 130% increase in VLP formation.

**Figure 3.**
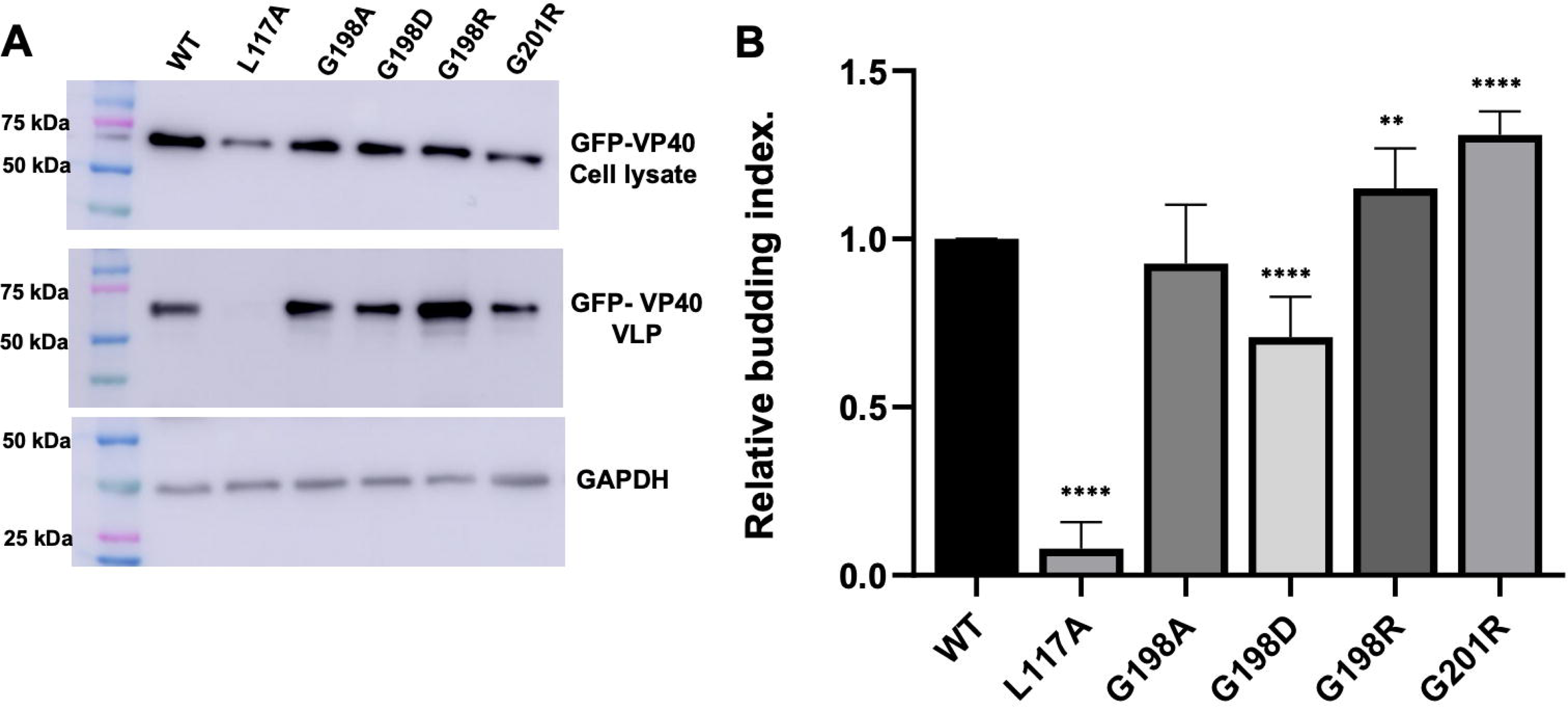
VLP analysis from VP40 and respective mutations expression in HEK293 cells. HEK293 cells were transfected with plasmid DNA encoding EGFP-WT-VP40, EGFP-VP40-L117A, EGFP-VP40-G198A, EGFP-VP40-G198D, EGFP-VP40-G198R, or EGFP-VP40-G201R. *A*, Representative blots of WT VP40 and respective mutations in the VLP or cell lysate fractions. The VLPs and cell lysates were collected 48 hrs after transfection and subjected to Western blot using anti-GFP. GAPDH was used as a loading control for these experiments. Molecular weight markers are shown for all blots to verify band size for EGFP-VP40 and GAPDH. *B*, ImageJ was used to quantify the band intensity for each sample n = 8 for WT-VP40 and G198R-VP40 and n = 4 for all other mutants. VLP budding efficiency was quantified and normalized to WT as 1.0. Error bars are the standard deviation; A Student’s t-test was performed compared with WT (**p<0.01 and ****p<0.0001).

### Molecular dynamic simulations of VP40 and cationic mutant mutations

To assess the effects of mutations on membrane binding we first performed molecular dynamic simulations of the WT dimer and arginine mutations (G198R and G201R) on VP40 membrane binding capabilities. An established bilayer system was used that recapitulates the asymmetric nature of the PM (35). Arginine mutations at positions 198 and 201 were introduced to the WT VP40 (PDB 4LDB), which resulted in appreciable changes to the overall positive nature of the CTD in this region (Fig. 4*A*). G198R and G201R also had increased H-bond contacts with the membrane compared to WT VP40 (Fig. 4*B*). A few increased membrane contacts were evident throughout the simulation time for G198R and G201R compared to WT VP40 (Fig. 5*A* and *B*, more detailed snapshots at 400 ns shown in Fig. S2). The mutations G198R and G201R lie in a flexible loop region, and as they interact with PM, preferentially with PI(4,5)P_2_, they also allow other residues in the loop 1 (219–233) and loop 2 (274–283) regions to reorient, resulting in more favorable PM interactions (Fig. 5).

**Figure 4.**
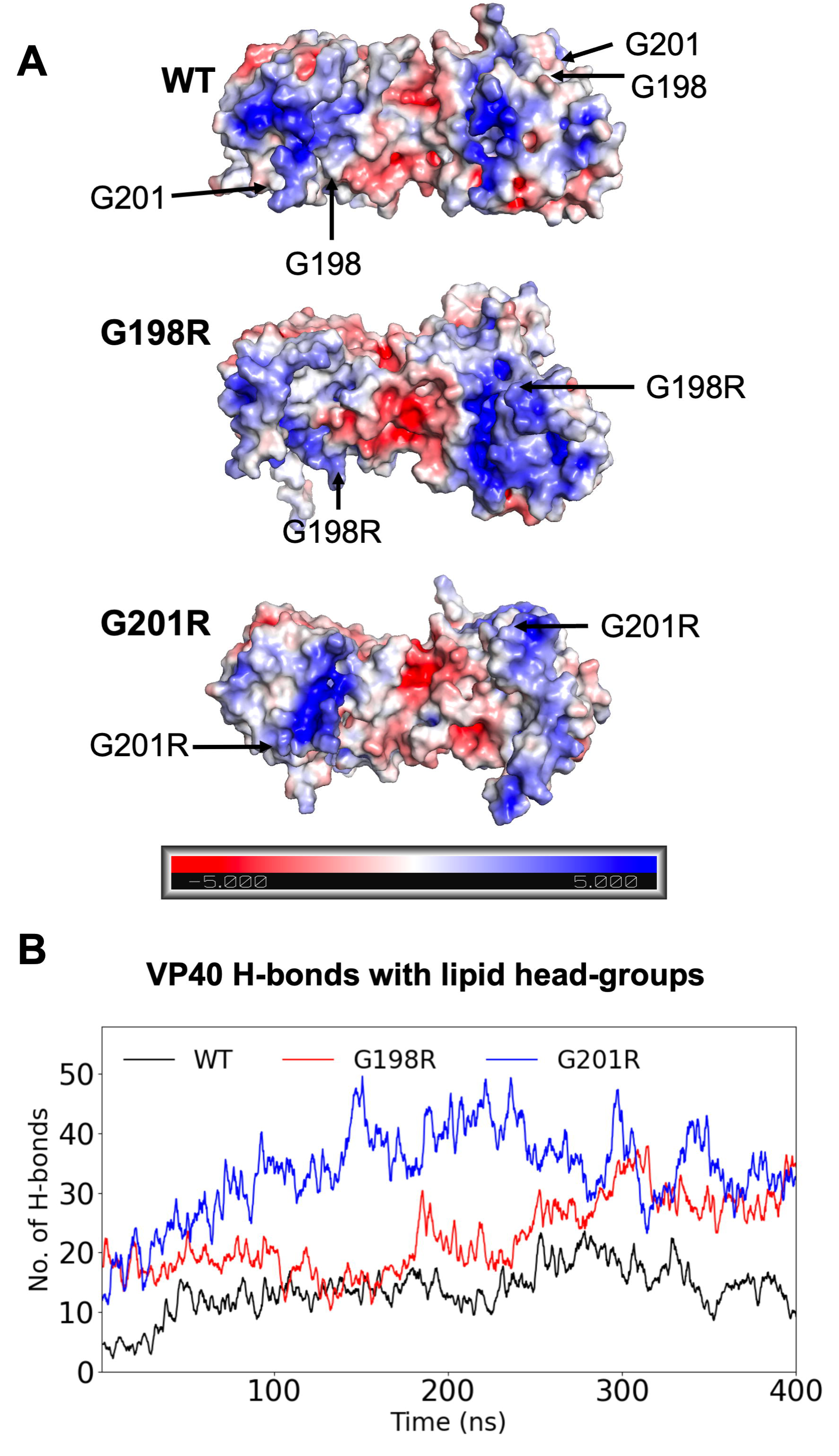
G198R and G201R have increased positive electrostatic charge on the CTD membrane-binding interface and make more H-bond contacts with membranes in MD simulations. *A*, The VP40 dimer structure is shown for WT, G198R and G201R revealing increased electrostatic charge on the membrane binding interface of the CTD near mutated glycine residues. *B*, Number of H-bonds with the lipid head groups made by the WT VP40 dimer as it associates with PM. The 400-ns simulation trajectories show that H-bonding is increased for G198R and G201R compared to WT.

**Figure 5.**
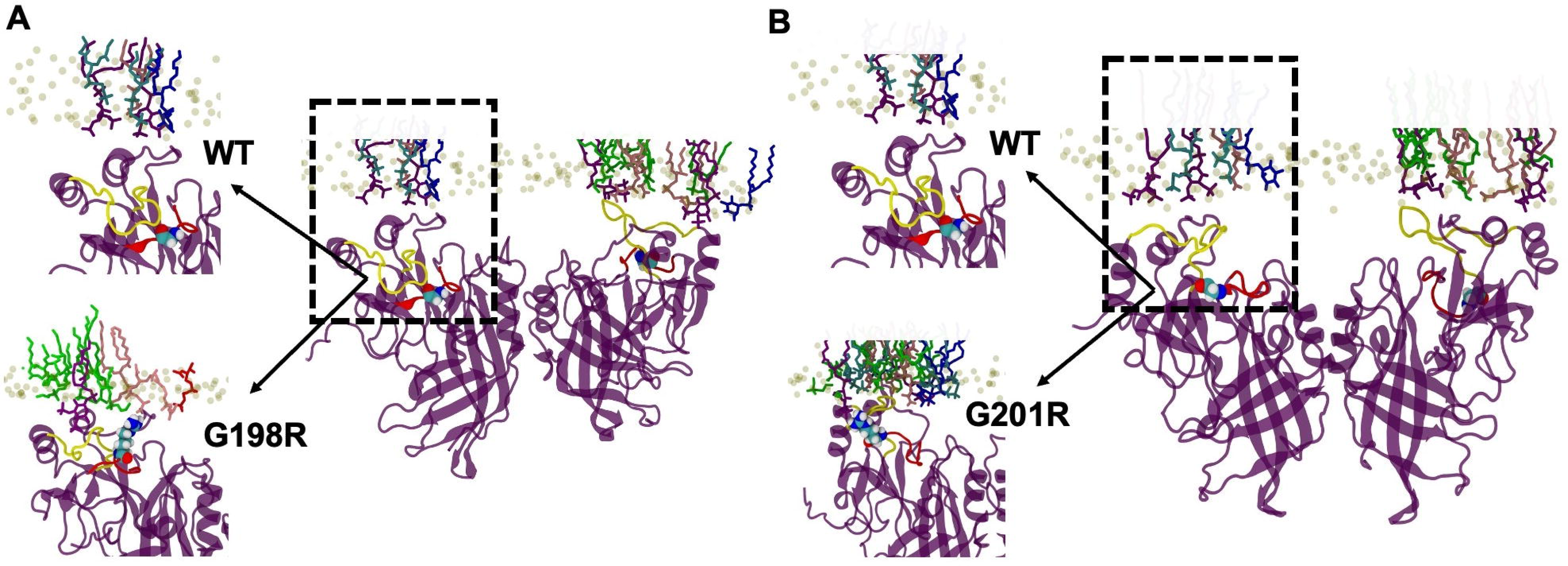
Snapshots of G198R and G201R molecular dynamics simulations reveal increased electrostatic interactions with the membrane compared to WT VP40. *A*, G198R increased membrane contacts between VP40 and the membrane interface compared to WT VP40 during the simulation. This includes increased contacts by R198, K221, K224, N227 and K279 residues. *B*, G201R is shown at 400 ns in contact with the model membranes. G201R leads to increased electrostatic association with the anionic membrane compared to WT as evidenced by increased interactions by residues K104, R201, K224, and S278.

### Introduction of cationic residues in VP40 increases affinity for PS and PI(4,5)P_2_ containing membranes

VP40 binds to PS and PI(4,5)P_2_ in the PM inner leaflet (10, 12), where CTD Lys residues are critical for lipid binding and VLP formation (8, 26, 33). Since mutations in the 198-GSNG-201 loop are located just beneath the membrane binding interface and those that increase positive charge led to an increase in PM assembly, budding, and increased H-bonding elucidated with MD simulations, we hypothesized that G198R and G201R were enhancing VP40 interactions with the anionic PM. To determine whether electrostatic changes at the membrane binding interface alter VP40 lipid binding ability, we evaluated the effects of G198A, G198R, G198D, and G201R on PS and PI(4,5)P_2_ binding using surface plasmon resonance (SPR) and a large multilamellar vesicle (MLV) binding assay, respectively. L117A was used as a control mutation as it inhibits dimer formation and subsequent PM localization and VLP formation.

WT VP40 and the mutations were expressed in *E. Coli* and purified by affinity chromatography. WT VP40 and LPI interface mutants (G198A, G198D, G198R and G201R) produced VP40 dimers and octamers in solution, so we separated them by size exclusion chromatography (Fig. S3). Even the G198D mutation, which had decreased PM localization and VLP production had a similar purification profile to WT, predominantly forming the VP40 dimer. This suggested G198D defects in PM localization and VLP budding are not caused by disruption of dimer formation. Since the dimers are the main building block for VP40 PM lipid interactions in EBOV assembly (8, 11) we only used the dimer in our lipid binding experiments (8). The L117A mutation produced little VP40 monomers and abundant octamers in solution (Fig. S3) as previously reported for the L117R (8, 26) and other VP40 dimer disrupting mutations (21). Currently there is no evidence to suggest that the octamer interacts significantly with the PM, as it has been shown to be important for the replication step of the viral lifecycle (9), Thus, we used the L117A monomer fraction to assess L117A binding affinity for PS.

To evaluate binding to PS in lipid vesicles, we prepared large unilammelar vesicles containing POPC/POPE/POPS (60:20:20) with a control flow cell of POPC/POPE (80:20). The SPR response unit signals versus protein concentration as well as the apparent *K*_d_ values for PS vesicle binding are shown in Fig. 6*A* and *B*, respectively. Mutations that increase the positive charge of VP40 (G198R and G201R) bind with ∼ 2-fold higher affinity for PS containing vesicles than WT VP40. In contrast, mutations that decrease the net charge (G198D) bind with ∼ 2-fold less affinity than WT VP40. This confirms our hypothesis from our cell studies that the changes in PM assembly and VLP budding are correlated with PM binding ability. The apparent *K*_d_ value for L117A monomer was difficult to estimate as the response at 2 μM was not sufficient to represent saturation of binding.

**Figure 6.**
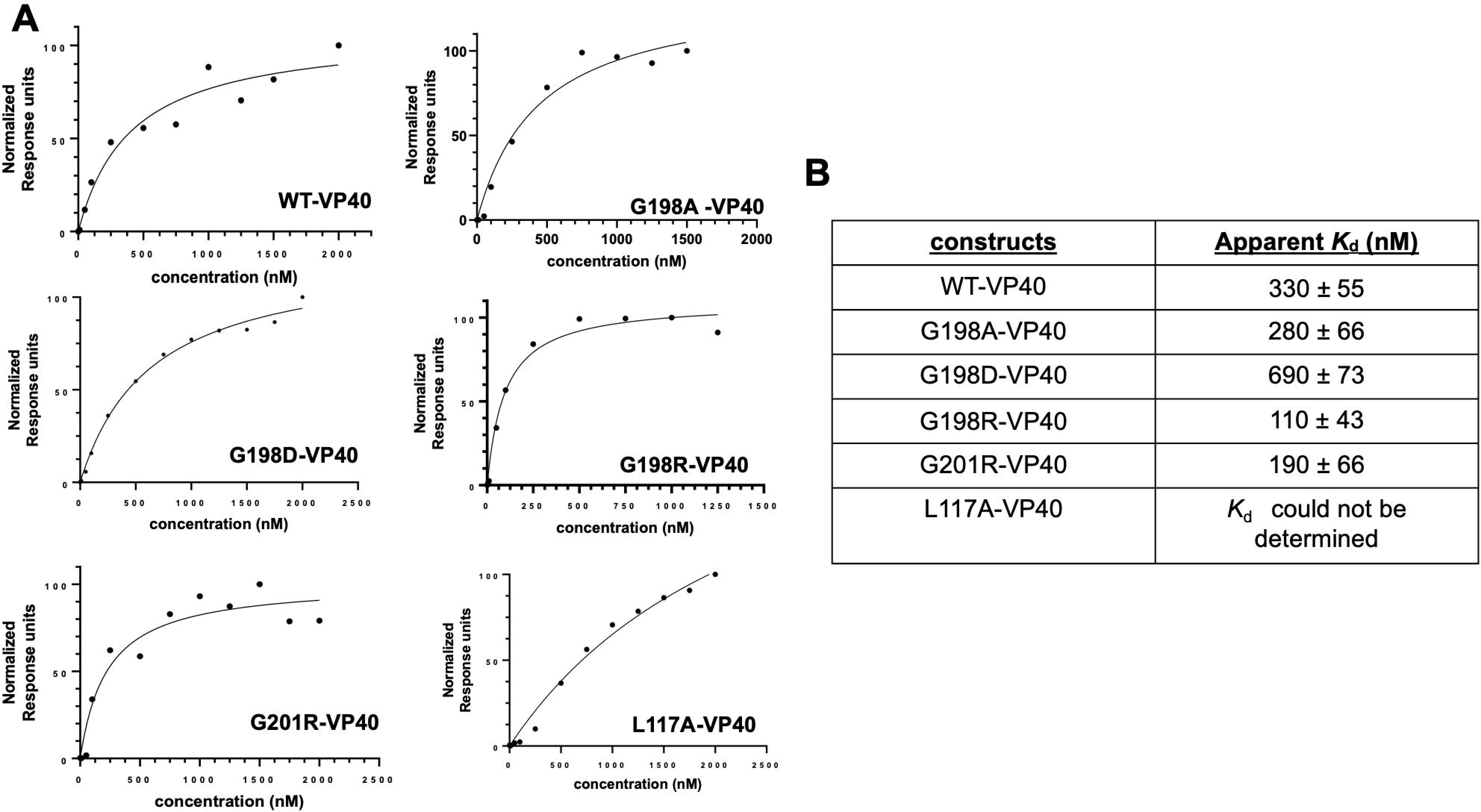
SPR binding measurements for WT VP40 and mutants for lipid vesicles containing POPS. *A*, Representative SPR binding curves for WT VP40 and mutants to LUVs (POPC:POPE:POPS (60:20:20)) containing 20% POPS, n = 4 for WT VP40 and n = 3 for all VP40 mutant measurements. *B*, Table showing the apparent binding affinities (*K*_d_) and STDEV of WT VP40 and mutants to lipid vesicles containing 20% POPS.

VP40 binding to PI(4,5)P_2_ was only quantified for select mutants (G198R and G201R) using a large multilamellar vesicle (MLV) pelleting assay. We monitored binding of the VP40 dimer to lipid vesicles containing POPC/POPE/PI(4,5)P_2_ (75:20:6.7). This assay allows for the quantitative comparison of how VP40 mutants affect the protein fraction that binds liposomes. Western blot analysis indicated the distribution of unbound protein in the supernatant and the fraction bound in the pellet (Fig 7*A*). Both the G198R and G201R mutations resulted in a significant increase in the fraction of VP40 bound to vesicles when compared to WT (Fig 7*B*).

**Figure 7.**
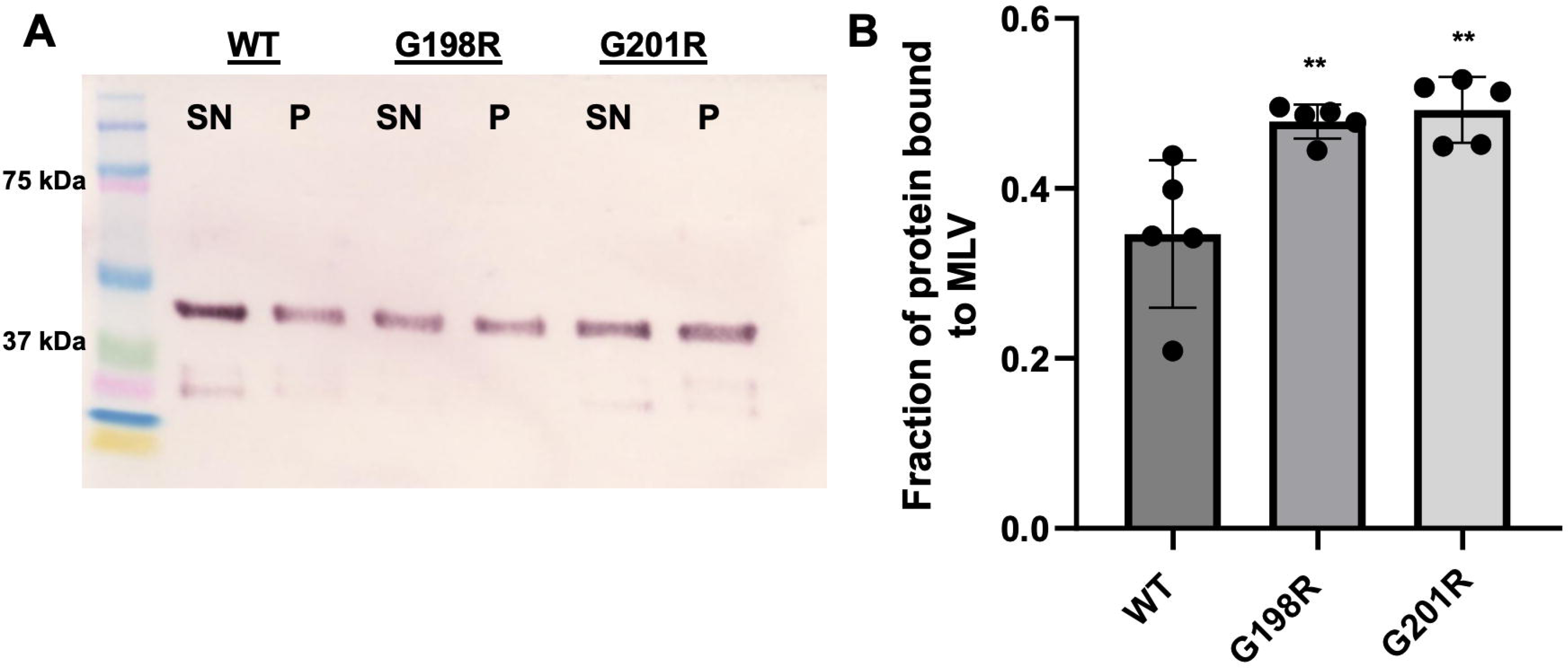
Assessment of WT VP40 and mutants association with lipid vesicles containing 6.7% PI(4,5)P_2_. *A*, Western blot of supernatant (SN) and pellet (P) fractions collected from the MLV sedimentation assays on VP40 constructs shown above the gels. MLVs (POPC:POPE: PI(4,5)P_2_ (67.3:20:6.7)) containing 6.7% PI(4,5)P_2_ were used. *B*, Quantification of MLV sedimentation assay results using Image J. The data represent the averages of five sedimentation assays, with the error bars representing the STDEV. A one-way ANOVA was performed with multiple comparisons compared with WT PM localization (**p < 0.01).

### VP40 CTD electrostatics influence PM assembly and VLP budding

The work presented thus far highlights the importance of VP40 electrostatic interactions on anionic lipid binding, PM assembly, and VLP formation in the EBOV viral lifecycle. It also underscores the sensitivity of the interaction with an additional charge of +1 having a profound effect on membrane binding and VLP formation. Previous research has shown that electrostatic interactions between the CTD of VP40 and membrane lipids are important for the initial association with the anionic membrane (10, 12, 26) and oligomerization of VP40 hexamers along the membrane (10, 12, 26, 33). Mutations that increased the charge difference between the membrane binding interface of VP40 (G198R and G201R) and the PM increased VP40 assembly and VLP budding while mutations that decrease this charge difference (G198D) reduced assembly and budding. To further evaluate how electrostatic changes on or near the VP40 CTD membrane binding interface altered VP40 assembly and budding, we evaluated the effects of mutating anionic residues exposed on the membrane binding surface of the VP40 CTD. This included aspartic acid and glutamic acid residues at this interface (Fig. 8) via assessment of the following mutations: D193A, D193K, E235A, E262A, and E265A.

**Figure 8.**
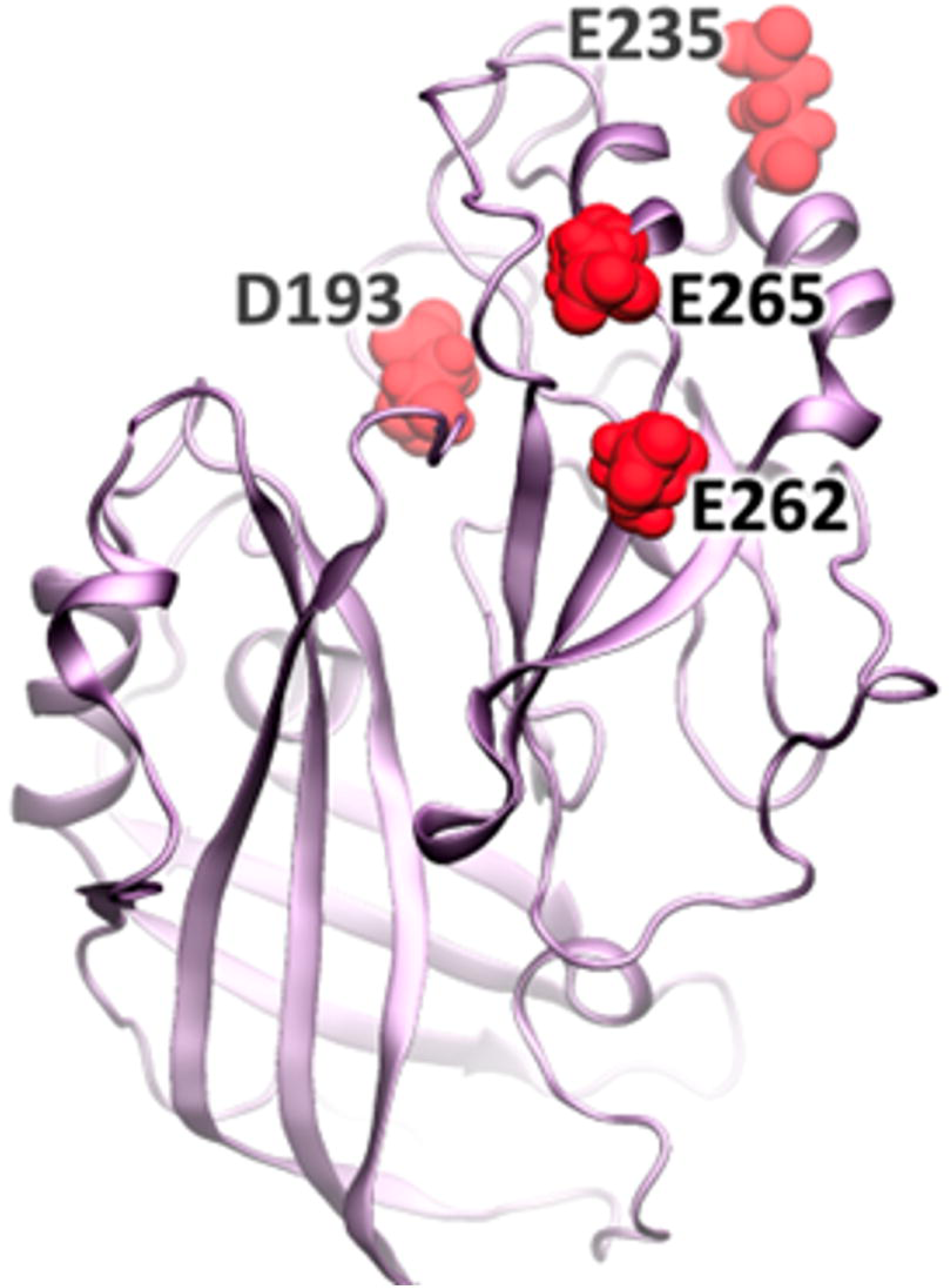
A schematic of a VP40 protomer displaying residues D193, E235, E262 and E265 at the CTD. The VP40 dimer structure (PDB ID: 4LDB) was used to model one of the protomers to highlight residues mutated to Ala or Lys. The ribbon diagram demonstrated the positioning of D193 and E235 directly at the lipid binding CTD interface with E262 and E265 are located near the CTD-CTD dimer interface.

We evaluated VP40 PM localization and VLP budding as described earlier. In a similar fashion to G198R and G201R, D193A, D193K, and E235A increased VP40 PM assembly (Fig. 9*A* and *B*) and VLP budding (Fig. 9*C* and *D*). Further, substitution of Asp with a Lys (D193K), an effective increase in positive charge by two, increased VP40 assembly and budding to a greater extent than the alanine substitution (D193A). E262A and E265A decreased PM assembly and VLP budding, in contrast to E235A, (Fig. 9*C* and *D*). Notably, Asp^193^ and Glu^235^ are located on the membrane binding interface of the VP40 CTD whereas Glu^262^ and Glu^265^ are located below the membrane binding interface and near the CTD-CTD dimer interface (Fig. 8). Hence, Glu^262^ or Glu^265^ do not likely interact directly with the membrane or greatly influence the charge of the membrane binding interface. We posit that the mutation of these residues affects dimer interactions that are necessary to stabilize the dimer at the CTD. Mutation would in turn destabilize the dimer leading to diminished PM assembly and VLP budding. These results further substantiate that an increase in positive charge at the VP40 CTD membrane binding interface increases VP40 PM binding and subsequent VP40 budding.

**Figure 9.**
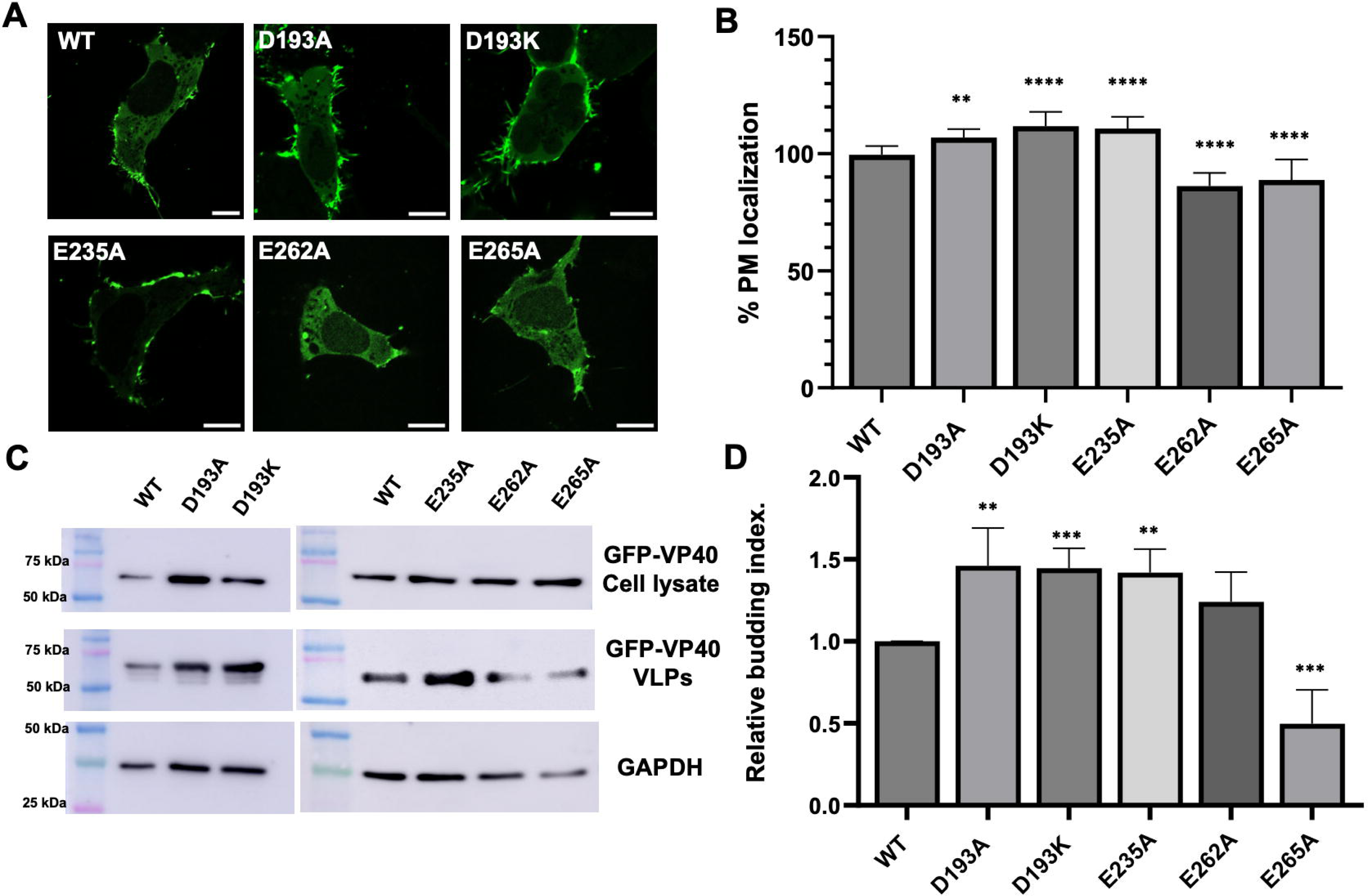
Cellular localization and VLP analysis of VP40 Asp and Glu mutations in HEK293 cells. *A*, Representative images of EGFP-WT-VP40, EGFP-VP40-D193A, EGFP-VP40-D193K, EGFP-VP40-E235A, EGFP-VP40-E262A, and EGFP-VP40-E265A in HEK293 cells 24 hours post-transfection. Scale bar = 10 μm. *B*, Average PM localization for each construct. Image analysis was done in Image J to determine percent PM localization (%PM localization) for each construct. %PM localization was defined as (PM intensity/(PM intensity+ cytosol intensity))*100. Data was normalized to %PM localization of WT as 100% localization. Error bars are the SEM from three independent experiments with at least 9 images analyzed per replicate. A one-way ANOVA was performed with multiple comparisons compared with WT PM localization (**p<0.01,***p<0.001 and ****p<0.0001). *C*, Representative blots of WT VP40 and respective mutations in the VLP or cell lysate fractions. The VLPs and cell lysates were collected 48 hrs after transfection and subjected to Western blot using anti-GFP. GAPDH was used as a loading control for these experiments. Molecular weight markers are shown for all blots to verify band size for EGFP-VP40 and GAPDH. *D*, ImageJ was used to quantify the band intensity for each sample n = 8 for WT-VP40 and n = 4 for all mutants. VLP budding efficiency was quantified and normalized to WT as 1.0. Error bars are the STDEV; A Student’s t-test was done compared with WT (**p<0.01 and ***p<0.001).

### EBOV VLP Entry is independent of VP40 CTD electrostatic charge

The co-expression of VP40 and the glycoprotein (GP) produces entry competent VLPs that can be used to assess VLP entry into cells (15, 18, 29, 35). To determine whether mutations at LPIs affect entry of VLPs, we performed a fluorescent-based entry assay using GFP-VLPs. HEK293 cells were incubated with 50 μg of VLPs derived from WT or mutant VP40 constructs. The L117A mutation and mock transfections were used as controls and did not have any detectable VLP entry (Fig. 10*A* and *B*). Although the VP40 mutations that were probed increased (e.g., G198R) or decreased (e.g., G198D) VLP formation based on electrostatic charge (Fig. 10), we wanted to compare equal levels of VLPs for the purposes of comparing viral entry. After incubation with GFP-VLPs, cells were stained with the Hoechst 33342 nuclear stain and fixed. Images were taken using confocal microscopy and the number of GFP particles present per cell nuclei were counted. In each case, mutations had no effect on VLP uptake. The mutations at the LPI (G198A, G198D, G198R and G201R) had no significant changes on VLP entry. This data suggests that even though mutations produce different amount of VLPs, mutations that increase VP40 overall positive charge increase VLP production while mutations that decrease VP40 overall positive charge decrease VLP production, the amount of VLP entry or uptake is similar. These findings in combination with the effects on assembly and lipid binding show that mutations like G198R affect the later stages of the viral lifecycle, leading to the production of more infectious particles available to infect new cells.

**Figure 10.**
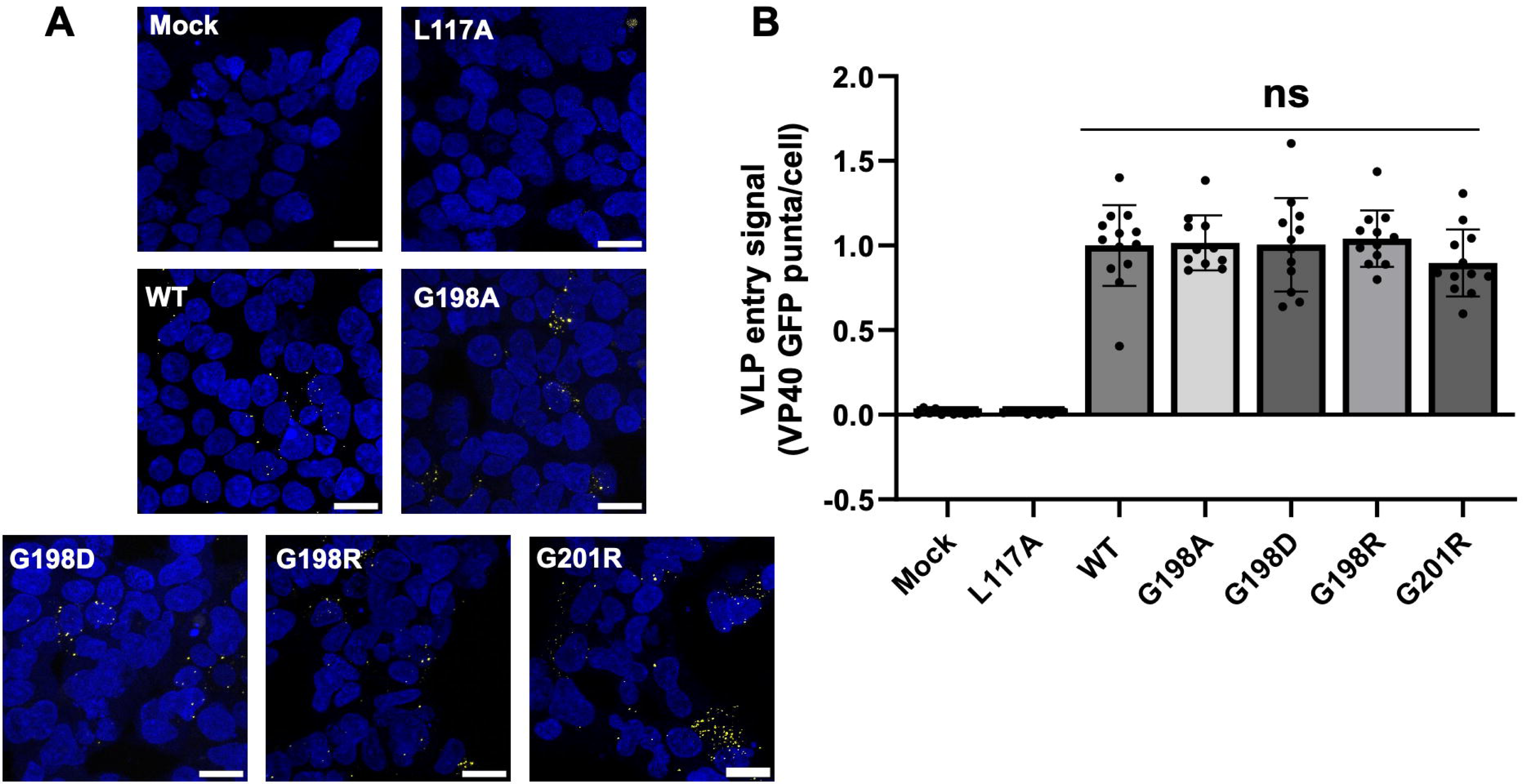
VLP entry assay using WT and mutant VP40 constructs in combination with GP. *A*, Representative images of EGFP-VLP entry comparing entry of VLPs produced from WT or mutant VP40 constructs into target HEK293 cells. A stack of 10-13 frames was acquired for each image and cells were stained with the Hoechst 33342 nuclear stain. Scale bar = 20 μm. VLPs (initially green) were recolored to yellow for easier observation in the figures. *B*, Quantification of VLP entry signal was performed by calculating the total number of VLPs (green puncta) / the total number of cells. The experiments were done in triplicate, N = 10 per replicate. Error bars represent STDEV. A one-way ANOVA was performed with multiple comparisons against WT.

## Discussion

In this study, we examined the critical role of electrostatic interactions at lipid-protein interaction interfaces between VP40 and the PM on viral assembly and budding. The investigation is primarily focused on elucidating the impact of mutations in the CTD region of VP40, particularly within the 198-GSNG-201 loop region on the protein’s interaction with anionic lipids. We used mutagenesis to evaluate how changing the electrostatic charge of VP40 at the lipid binding interface affects different stages of the viral lifecycle. Confocal imaging and assessment of VLP budding showed that the electrostatic charge in this loop region affects VP40 assembly and VLP budding efficiency. Mutations leading to an increase in the positive charge of the protein, such as G198R and G201R, enhanced VP40 PM assembly and VLP budding. Conversely, mutations reducing the overall charge, exemplified by G198D, displayed compromised assembly and budding efficiency. The observed effects of these mutations primarily impact the later stages of the viral lifecycle, particularly in the efficiency of PM membrane association and VLP budding but have no effect on viral entry.

WT VP40 can conformationally rearrange into the dimer for further oligomerization and the RNA-binding octameric ring (8, 19, 21). Previous work has shown that mutations can alter VP40 assembly and budding by changing the distribution of the dimer and octamer formed (8, 21, 26). Our assessment of VP40 oligomerization *in vitro* showed no difference in detected VP40 oligomers in their purification profiles. This is particularly interesting because it shows that the reduction in assembly and budding by the G198D mutant is not due to dimer formation or trafficking defects. To confirm that effects observed were directly correlated to the lipid-protein interaction interface, we compared the WT to VP40 construct bearing an L117A mutation located at the NTD-NTD dimer interface. Similarly, to the previously studied L117R mutation (8, 26), L117A does not localize to the PM in cells and remains diffusely distributed throughout the cell. Further, L117A doesn’t form detectable VLPs. The L117A mutation however predominantly formed the VP40 octamer and a small fraction of the monomer. This further supports that different VP40 interfaces play different roles and are involved in different stages of the lifecycle. The CTD is important for VP40 interactions with the membrane affecting later stages of the viral lifecycle while the NTD important for dimer formation facilitating the role of VP40 in inhibiting viral genome replication and transcription (9, 24, 36). Further, VP40 dimer stability is maintained by a different NTD interface than that of the NTD in the VP40 octamer, where dimer structure is critical for proper trafficking of VP40 to the PM (8).

VP40 selectively binds to anionic lipids, PS and PI(4,5)P_2_ in particular in the inner leaflet of the PM. Here the VP40 dimer interacts with the PM through key lysine residues (especially Lys^221^, Lys^224^, Lys^225^, Lys^274^, and Lys^275^) in its C-terminal domain (CTD) (8, 26, 37). These basic residues interact electrostatically with the PM to facilitate matrix filament assembly. The 198-GNSG-201 loop region is adjacent to the membrane interacting loop residues, Lys^224^ and Lys^225^, suggesting that it may play a critical role in VP40 lipid binding. MD simulations suggested that arginine mutations (G198R and G201R) interact with more membrane lipids and will form more H-bonds bonds with membrane lipids. *In vitro* binding studies with PS and PI(4,5)P_2_ confirmed this hypothesis, where the G198R and G201R mutations increased the affinity of VP40 to PS and PI(4,5)P_2_ containing vesicles. In contrast, G198D decreased affinity for PS containing vesicles by more than ∼2 fold in SPR binding experiments. In addition to increasing the affinity for anionic lipids, MD simulations show that arginine substitutions lead to a conformational change in the 198-GSNG-201 loop exposing more residues to interact with the membrane. The importance of electrostatic interactions on VP40 assembly at the PM and budding have been highlighted in previous studies (26). Here we demonstrated the sensitivity of these interactions by the significant effects observed by altering VP40 overall charge density by ∼ +1 or -1. The study’s findings emphasize the potential therapeutic implications of manipulating electrostatic interactions at lipid-protein interfaces to inhibit viral assembly and subsequent infectivity.

Furthermore, the study demonstrates the significance of the overall surface charge of the VP40 lipid binding interaction interface outside of the 198-GSNG-201 motif. This is evidenced by increasing VP40 electrostatic charge by mutating anionic residues (Asp and Glu) in the CTD. Mutations that increased the electrostatic charge of the VP40 surface in regions interacting directly with the membrane (e.g., D193A, D193K, and E235A) similarly led to enhanced assembly and budding. However, it was noted that mutations targeting residues located at the CTD but not directly interacting with the membrane (E262A and E265A), did not yield a similar effect on assembly and budding, reinforcing the specificity of these interactions in driving the viral assembly process. Since Glu^262^ and Glu^265^ are in the CTD-CTD dimer interface we hypothesize that they play a role in stabilizing the dimer or dimer-dimer interactions. Hence, mutations at these residues possibly augment important interactions leading to the destabilization of the dimer. This may affect the population of dimer that can interact with the membrane.

This investigation underscores the crucial role of electrostatic interactions between specific CTD regions of VP40 and anionic lipids in the PM, providing valuable insights into the molecular mechanisms governing EBOV assembly and budding. Understanding these interactions may offer potential avenues for the development of targeted antiviral strategies aimed at disrupting viral assembly and inhibiting the spread of EBOV infections. This study may also help springboard understanding of mutations of VP40 that have occurred in previous EBOV epidemics in samples sequenced from patients (www.nextstrain.org). A wide spectrum of mutations has been discovered in both the NTD and CTD. Several VP40 mutations discovered are of immediate relevance to this study as they alter the electrostatic nature of the VP40 CTD membrane binding interface. These mutations include G223R (2018-2021 Ebola epidemic BEN42629), G226R (2013-2016 West African Ebola epidemic LIBR10224), Q245R (2013-2016 West African Ebola epidemic EM_COY_2015_016267), H269R (2013-2016 West African Ebola epidemic EM_076615 and 2018-2021 Ebola epidemic BTB7117), and D312G (2018-2021 Ebola epidemic BTB238). G223R and G226R are of note as they lie in a loop region previously found to interact with PS and PI(4,5)P_2_ containing membranes through Lys^224^ and Lys^225^. Introduction of an arginine in this loop may further enhance electrostatic association of the PM inner leaflet. Q245R introduces a positive charge in the CTD domain near where dimer CTDs interact with one another. H269R introduces additional positive charge (as His can be protonated below physiological pH) adjacent to critical PS and PI(4,5)P_2_ binding residues Lys^270^, Lys^274^, and Lys^279^. Meanwhile, D312G reduces anionic charge just below the membrane binding interface. In consideration of this study, it is tempting to hypothesize these mutations will increase PM interactions and virus assembly. However, how these mutations contribute to VP40 structure and function properties will require *in vitro* and cellular analysis of VP40 lipid interactions building off the current studies.

### Experimental procedures

#### Plasmids

The mammalian expression plasmid for VP40 was created by subcloning VP40 into enhanced green fluorescent protein (EGFP)-pcDNA3.1 as previously described (27). EGFP-VP40 mutants in pcDNA3.1 were prepared and purchased from Gene Universal (Newark, DE). The His6-eVP40-pET46 expression vector was a kind gift from Dr. Erica Ollmann Saphire (La Jolla Institute for Immunology). The His6-eVP40-pET46 mutants were also prepared and purchased from Gene Universal.

#### Cell culture and imaging

Human embryonic kidney (HEK293) cells were cultured and maintained at 37°C in a 5% CO_2_ humidified incubator supplemented with Dulbecco’s Modified Eagle’s Medium (low glucose) (Thermo Fisher Scientific, Waltham, MA) containing 10% fetal bovine serum (Nucleus Biologics, San Diego, CA) and 1% Pen/Strep (Thermo Fisher Scientific, Waltham, MA). After trypsinization, cells were transferred to a 35 mm glass bottom dish (MatTek, Ashland, MA). Cells were then grown to 70–80% confluency and transfected with 2.5 μg DNA/dish using Lipofectamine^TM^ 2000 (Invitrogen, Carlsbad, CA) according to the manufacturer’s protocol. Transfections were carried out for 8 hrs to assess the effects of VP40 mutations at an early time point (e.g., low level of expression) and for 24 hrs to assess the effects of these mutations at a later time point. Cells were stained with the Hoechst 33342 nuclear stain and the WGA-Alexa647 membrane stain (Thermo Fisher Scientific) and imaged using a Nikon A1R confocal microscope with a plan apochromat 60x 1.4 NA oil objective. The 488 nm laser line was used for the excitation of EGFP, the 405 nm line was used for the excitation of the Hoechst nuclear stain and the 640 nm line was used for the excitation of the WGA-Alexa647 PM stain. PM localization was calculated as follows: (PM intensity/(PM intensity +cytosol intensity))*100. WT EGFP-VP40 and EGFP were used to normalize the PM localization signal to 100% and 0%, respectively, to assess several VP40 mutations for PM localization. At least 24 cells were imaged for WT, EGFP, and each mutation over three independent experiments.

#### Virus-like particle collection

HEK293 cells were grown in a 100 mm round dish and transfected with 15 μg of DNA as previously described (12) .VLPs were collected 48 hrs post transfection and budding assays were performed as described previously (12). In brief, VLP containing supernatants were harvested from cells and clarified through low-speed centrifugation (1000 x *g* for 10min). Clarified VLPs were loaded onto a 20% sucrose cushion in STE buffer (10□mM TRIS, pH 7.6, containing 100□mM NaCl and 1□mM EDTA), isolated through ultracentrifugation (110,000 x *g* for 3 hrs), and resuspended in STE buffer. Cell lysate samples were harvested and lysed on ice with RIPA buffer (50 mM TRIS, pH 7.4, containing 150□mM NaCl, 5□mM EDTA pH□8, 1% Triton X-100, 0.1% SDS, and 0.5% deoxycholic acid) supplemented with 1% Halt™ Protease Inhibitor Cocktail (Thermo Fisher Scientific). Samples were stored at -20°C for western blot analysis.

#### Western blot analysis

Proteins from cell lysates and VLPs were separated by size using 10% SDS-PAGE. Following transfer onto a nitrocellulose membrane, the nitrocellulose membranes were blocked with 5% milk in TBS-T (20 mM TRIS, pH 8.0, containing 150 mM NaCl and 0.1% Tween) and analyzed with their respective antibodies. GFP-VP40 was detected using the anti-GFP HRP antibody (Abcam, Cambridge, United Kingdom). GAPDH was detected using the mouse anti-GAPDH primary antibody (MilliporeSigma^TM^ Burlington, MA) and the sheep anti-mouse-HRP secondary antibody (Abcam). Antibodies were detected using an enhanced chemiluminescence (ECL) detection reagent (Bio-Rad, Hercules, CA) and imaged on an Amersham Imager 600. All quantitative analysis were performed using densitometry analysis in ImageJ. Following ECL detection, VP40 cell lysate (VP40_CL_) expression was normalized to the respective GAPDH band density. The relative budding index was calculated according to the ratio of density_VLP_/density_CL+VLP_ (where density_VLP_ is the VP40 VLP band density and density_CL+VLP_ is the VP40 cell lysate + VLP band density).

#### Molecular Dynamics Simulations

The VP40 dimer structure was obtained from the Protein Data Bank (PDB ID: 4ldb) (8, 38) and missing residues were added with Modeller (39). The protein-PM system was built using the Charmm-Gui (40) membrane builder web interface (41). The PM consists of cholesterol (CHOL), phosphatidylcholine (POPC), phosphatidylethanolamine (POPE), phosphatidylinositol (POPI), phosphatidylinositol 4,5-bisphosphate (PIP_2_), phosphatidylserine (POPS), and palmitoyl sphingomyelin (PSM) molecules (39, 42, 43). The resulting system contained 226 lipids in the outer leaflet and 227 lipids in the inner leaflet, with CHOL:POPC:POPE:POPS:POPI: PIP_2_:PSM ratio of 21:11:31:17:3:7:10. The protein-PM system was solvated with TIP3 water molecules and 0.15M KCl in a cubic box. The G198R and G201R mutations were introduced and systems were setup with Charmm-Gui (40, 44). The total number of atoms in the final solvated systems (protein, membrane, water, and ions) was 197,791 atoms for the WT, 190,180 for the G198R, and 186,934 for the G201R system.

Molecular dynamics simulations were performed with NAMD 3.0 (45) using CHARMM36 force field (46). The particle mesh Ewald (PME) method (47) was used for electrostatic interactions and the covalent bonds involving the hydrogen atoms were restrained with the SHAKE (48) algorithm. For each system, a 10,000-step minimization followed by a 6-step equilibration run was performed before starting the NVT production run at 303.15 K with a timestep of 2 fs. The pressure and temperature were regulated by the Nose– Hoover Langevin-piston method (49) and the Langevin coupling, respectively. Visual Molecular Dynamics (VMD) 1.9.4 (50) was used to analyze the simulation trajectories and to render images

#### Protein purification

WT His_6_-VP40 and mutants were grown and purified as previously described (12). Following Ni-NTA affinity chromatography elution, WT VP40 and VP40 mutants were further purified using size exclusion column on a Hiload 16/600 Superdex 200 pg column (Cytiva Life Sciences, Marlborough, MA). The desired fractions were collected, concentrated, and stored in 10 mM HEPES, pH 8.0, containing 150 mM NaCl. A sample of the purified protein was run on an SDS-PAGE gel and stained with Coomassie Brilliant Blue to confirm the appearance of a strong band at ∼40 kDa. Protein concentrations were determined using the Pierce^TM^ BCA Protein Assay (Thermo Fisher Scientific) and the protein was stored at 4°C for no longer than 2 weeks.

#### Lipids and LUV preparation

POPC (#850457), POPE (#850757), POPS (#840034) and PI(4,5)P_2_ (#850155) were purchased from

Avanti Polar Lipids, Inc. (Alabaster, AL) and stored in chloroform and/or methanol at -20°C until use. For large unilamellar vesicle (LUV) preparation, lipid mixtures were prepared at the indicated compositions (see surface plasmon resonance section), dried down to lipid films under a continuous stream of N_2_, and stored at -20°C until further use. On each day of experiments, LUVs were brought to room temperature, hydrated in SPR buffer (10□mM HEPES, pH 7.4 containing 150 mM NaCl), vortexed vigorously, and extruded through a 100□nm filter 17 times.

#### Surface plasmon resonance

To determine the affinity of WT His_6_-VP40 WT and mutants to LUVs with PS, SPR was performed. SPR experiments were performed at 25°C using a Biacore X100 as described previously (10, 51). In brief, an L1 chip was coated at 5□μl/min with LUVs containing 0% PS (POPC:POPE (80:20)) on flow cell 1 and 20% mol PS or on flow cell 2 (PS was added at the expense of POPC). The LUV conjugated chip was stabilized by washing with 50 □mM NaOH and blocked with 0.1□mg/ml BSA (in SPR buffer) at a flow rate of 10□μl/min. For quantitative affinity analysis, each concentration of VP40 was injected for 510□s at a flow rate of 10□μl/min with a 180□s delay, and the difference in response between flow cell 1 and flow cell 2 was recorded (ΔRU). The apparent *K*_d_ of vesicle binding was determined using the non-linear least squares analysis: R_eq_ = R_max_/(1+*K*_d_/C) where *R*_eq_ (measured in RU) is plotted against protein concentration (C). *R*_max_ is the□theoretical maximum RU response and *K*_d_ is the apparent membrane affinity. Data were fit using the Kaleidagraph fit parameter of□(m0*m1)/(m0□+□m2); m1□=□1, 100; m2□=□1. ΔRU data were normalized in GraphPad Prism 8 for windows (La Jolla, CA) and plotted in Kaleidagraph (Reading, PA).

#### Multilamellar (MLV) sedimentation assay

MLV sedimentation assays were performed as previously described (12). In brief, an MLV emulsion containing 6.7% mol PI(4,5)P_2_ was made and dried to a film under nitrogen. The film was then hydrated in VP40 storage buffer (10 mM HEPES, pH 8.0, containing 150 mM NaCl) to a lipid concentration of 2.0 mM and incubated for 10 minutes at 37°C. Lipids were then vortexed vigorously for 60 seconds to form MLVs. Lipid suspension and protein was combined to a final lipid concentration of 1 mM and final protein concentration of 1 μM per sample. The mixture was then incubated at room temperature for 20 minutes. The supernatant (soluble protein) and pellet (lipid associated protein) were separated using ultracentrifugation (75,000 x *g* for 20 minutes). The pellet was resuspended in VP40 storage buffer to the same volume as the supernatant and vortexed to resuspend the pellet. Equal volumes of supernatant and pellet samples were resolved using 10% SDS-PAGE. Following transfer onto a nitrocellulose membrane the proteins were detected using the Anti-poly Histidine−Alkaline Phosphatase (AP) antibody (MilliporeSigma^TM^ Burlington, MA). Antibodies were detected using a colorimetric AP substrate reagent (Bio-Rad, Hercules, CA) and imaged on an Amersham Imager 600. All quantitative analyses were performed using densitometry analysis in ImageJ. To calculate the fraction of protein bound the following equation was used: density_pellet_/(density_supernatant_ +density_pellet_).

#### EGFP-VP40 entry assay

VLPs produced from HEK293 cells expressing GFP-VP40 and GP were purified as described in the VLP collection section. GFP-VP40 entry assays were performed as described previously (52, 53). In brief, following ultra-centrifugation, VLPs were resuspended in STE buffer and protein concentrations in VLPs were determined using the Pierce BCA assay. HEK293 cells were transfected with TIM-1 for 24□h prior to incubation with VLPs. 50 μg GFP-VP40 VLPs were added to TIM-1 expressing HEK293 cells, incubated 4°C for 1 h and allowed to incubate for 1□h at 37°C. Cells were then stained with the Hoechst 33342 nuclei stain, rinsed with PBS, fixed with 4% paraformaldehyde in PBS, and stored at 4°C until imaging. During image acquisition, z-stacks were acquired of 10–15 frames (1□μm steps). Quantification of VLP entry signal was performed by calculating the total number of VLPs (green puncta)/the total number of cells. The presence of EGFP-VP40 and GP was detected using western blot analysis as previously described (see Western blot analysis above). GP was detected using the Rabbit anti-EBOV GP primary antibody (IBT Bioservices, Rockville, MD) and the Goat anti-Rabbit secondary antibody (Abcam).

## Data Availability

The reported data within the article will be shared by the corresponding author upon request.

### Supporting Information

This article contains supporting information.

### Author contributions

Conceptualization: B.B.M. and R.V.S.; Data curation: B.B.M., T.S., and P.P.C.; Formal analysis: B.B.M., T.S., and P.P.C.; Funding acquisition: P.P.C. and R.V.S.; Investigation: B.B.M. and T.S.; Methodology: B.B.M., T.S., and P.P.C.; Project administration: P.P.C. and R.V.S.; Resources: P.P.C. and R.V.S.; Supervision:

P.P.C. and R.V.S.; Writing – original draft: B.B.M.; Writing – review & editing: B.B.M., T.S., P.P.C., and R.V.S.

### Funding and additional information

This work was supported by a National Institutes of Health grant to R.V.S. (AI158220).

## Abbreviations

CTD: C-terminal domain
EBOV: Ebola virus
NTD: N-terminal domain
PI(4,5)P_2_: phosphatidylinositol-(4,5)-bisphosphate
PM: plasma membrane
PS: phosphatidylserine
VP40: viral protein 40 kDa

## References

1. Carlson, C. J., Boyce, M. R., Dunne, M., Graeden, E., Lin, J., Abdellatif, Y. O., et al. The World Health Organization’s disease outbreak news: A retrospective database. (2023) PLoS Glob. Public Health 3, e0001083

2. Almeida-Pinto, F., Pinto, R., and Rocha, J. (2024) Navigating the complex landscape of ebola infection treatment: A review of emerging pharmacological approaches. Infect. Dis. Ther. 13, 21–55

3. Rijal, P. and Donnellan, F. R. (2023) A review of broadly protective monoclonal antibodies to treat Ebola virus disease. Curr. Opin. Virol. 61, 101339

4. Wolf, J., Jannat, R., Dubey, S., Troth, S., Onorato, M. T., Coller, B.A., et al. (2021) Development of pandemic vaccines: ERVEBO case study. Vaccines 9, 190

5. Stahelin, R. V. (2014) Could the Ebola virus matrix protein VP40 be a drug target? Expert Opin. Ther. Targ. 18, 115–120

6. Jain, S., Martynova, E., Rizvanov, A., Khaiboullina, S., and Baranwal, M. (2021) Structural and functional aspects of Ebola virus proteins. Pathogens 10, 1330

7. Harty, R. N. (2009) No exit: Targeting the budding process to inhibit filovirus replication. Antivir. Res. 81, 189–197

8. Bornholdt, Z. A., Noda, T., Abelson, D. M., Halfmann, P., Wood, M. R., Kawaoka, Y. et al. (2013) Structural rearrangement of Ebola virus VP40 begets multiple functions in the virus life cycle. Cell 154, 763–774

9. Hoenen, T., Volchkov, V., Kolesnikova, L., Mittler, E., Timmins, J., Ottmann, M. et al. (2005) VP40 Octamers Are Essential for Ebola Virus Replication *J*. Virol. 79, 1898–1905

10. Adu-Gyamfi, E., Johnson, K. A., Fraser, M. E., Scott, J. L., Soni, S. P., Jones, K. R. et al. (2015) Host cell plasma membrane phosphatidylserine regulates the assembly and budding of Ebola virus. J. Virol. 89, 9440–9453

11. Wan, W., Clarke, M., Norris, M. J., Kolesnikova, L., Koehler, A., Bornholdt, Z. A. et al. (2020) Ebola and Marburg virus matrix layers are locally ordered assemblies of VP40 dimers. eLife 9, e59225

12. Johnson, K. A., Taghon, G. J. F., Scott, J. L., and Stahelin, R. V. (2016) The Ebola Virus matrix protein, VP40, requires phosphatidylinositol 4,5-bisphosphate (PI(4,5)P2) for extensive oligomerization at the plasma membrane and viral egress. Sci. Rep. 6, 19125–19125

13. Jasenosky, L. D., Neumann, G., Lukashevich, I., and Kawaoka, Y. (2001) Ebola virus VP40-induced particle formation and association with the lipid bilayer. J. Virol. 75, 5205–5214

14. Stahelin, R. V. (2014) Membrane binding and bending in Ebola VP40 assembly and egress. Front. Microbiol. 5, 300–300

15. Noda, T., Sagara, H., Suzuki, E., Takada, A., Kida, H., and Kawaoka, Y. (2002) Ebola virus VP40 drives the formation of virus-like filamentous particles along with GP. J. Virol. 76, 4855–4865

16. Noda, T., Ebihara, H., Muramoto, Y., Fujii, K., Takada, A., Sagara, H. et al. (2006) Assembly and budding of Ebolavirus. PLoS Pathog. 2, e99–e99

17. Johnson, R. F., Bell, P., and Harty, R. N. (2006) Effect of Ebola virus proteins GP, NP and VP35 on VP40 VLP morphology Virol. J. 3, 31

18. Saeed, M. F., Kolokoltsov, A. A., Albrecht, T., and Davey, R. A. (2010) Cellular entry of ebola virus involves uptake by a macropinocytosis-like mechanism and subsequent trafficking through early and late endosomes PLoS Pathog. 6, e1001110

19. Dessen, A., Volchkov, V., Dolnik, O., Klenk, H.-D., and Weissenhorn, W. (2000) Crystal structure of the matrix protein VP40 from Ebola virus. EMBO J. 19, 4228–4236

20. Adu-Gyamfi, E., Soni, S. P., Jee, C. S., Digman, M. A., Gratton, E., and Stahelin, R. V. (2014) A loop region in the N-terminal domain of Ebola virus VP40 is important in viral assembly, budding, and egress. Viruses 6, 3837–3854

21. Narkhede, Y. B., Bhardwaj, A., Motsa, B. B., Saxena, R., Sharma, T., Chapagain, P. P. et al. (2023) Elucidating residue-level determinants affecting dimerization of Ebola virus matrix protein using high-throughput site saturation mutagenesis and biophysical approaches. J. Phys. Chem. B 127, 6449–6461

22. Yamayoshi, S., Noda, T., Ebihara, H., Goto, H., Morikawa, Y., Lukashevich, I. S. et al. (2008) Ebola virus matrix protein VP40 uses the COPII transport system for its intracellular transport. Cell Host Microbe 3, 168–177

23. Bhattarai, N., Pavadai, E., Pokhrel, R., Baral, P., Hossen, M. L., Stahelin, R. V. et al. (2022) Ebola virus protein VP40 binding to Sec24c for transport to the plasma membrane. Proteins 90, 340–350 10.1002/prot.26221

24. Soni, Smita P., Adu-Gyamfi, E., Yong, Sylvia S., Jee, Clara S., and Stahelin, R. V. (2013) The Ebola virus matrix protein deeply penetrates the plasma membrane: An important step in viral egress. Biophys. J. 104, 1940–1949

25. Amiar, S., and Stahelin, R. V. (2020) The Ebola virus matrix protein VP40 hijacks the host plasma membrane to form virus envelope *J*. Lipid Res. 61, 971–971

26. Del Vecchio, K., Frick, C. T., Gc, J. B., Oda, S.-I., Gerstman, B. S., Saphire, E. O. et al. (2018) A cationic, C-terminal patch and structural rearrangements in Ebola virus matrix VP40 protein control its interactions with phosphatidylserine. J. Biol. Chem. 293, 3335–3349

27. Adu-Gyamfi, E., Digman, M. A., Gratton, E., and Stahelin, R. V. (2012) Investigation of Ebola VP40 assembly and oligomerization in live cells using number and brightness analysis. Biophys. J. 102, 2517–2525

28. Adu-Gyamfi, E., Soni, S. P., Xue, Y., Digman, M. A., Gratton, E., and Stahelin, R. V. (2013) The Ebola virus matrix protein penetrates into the plasma membrane: a key step in viral protein 40 (VP40) oligomerization and viral egress. J. Biol. Chem. 288, 5779–5789

29. Husby, M. L., Amiar, S., Prugar, L. I., David, E. A., Plescia, C. B., Huie, K. E., et al. (2022) Phosphatidylserine clustering by the Ebola virus matrix protein is a critical step in viral budding. EMBO Reports 23, e51709

30. Scianimanico, S., Schoehn, G., Timmins, J., Ruigrok, R. H. W., Klenk, H.-D., and Weissenhorn, W. (2000) Membrane association induces a conformational change in the Ebola virus matrix protein. EMBO J. 19, 6732–6741

31. Ruigrok, R. W. H., Schoehn, G., Dessen, A., Forest, E., Volchkov, V., Dolnik, O. et al. (2000) Structural characterization and membrane binding properties of the matrix protein VP40 of ebola virus. J. Mol. Biol. 300, 103–112

32. Soni, S. P., and Stahelin, R. V. (2014) The Ebola virus matrix protein VP40 selectively induces vesiculation from phosphatidylserine-enriched membranes. J. Biol. Chem. 289, 33590–33597

33. Johnson, K. A., Budicini, M. R., Bhattarai, N., Sharma, T., Urata, S., Gerstman, B. S., et al. (2024) PI(4,5)P2 Binding Sites in the Ebola Virus Matrix Protein Modulate Assembly and Budding J. Lipid. Res. articles in press, 100512

34. Johnson, K. A., Bhattarai, N., Budicini, M. R., LaBonia, C. M., Baker, S. C. B., Gerstman, B. S. et al. (2021) Cysteine mutations in the Ebolavirus matrix protein VP40 promote phosphatidylserine binding by increasing the flexibility of a lipid-binding loop. Viruses 13, 1375

35. Gc, J. B., Gerstman, B. S., and Chapagain, P. P. (2017) Membrane association and localization dynamics of the Ebola virus matrix protein VP40. Biochim Biophys Acta Biomembr 1859, 2012–2020

36. Manicassamy, B., and Rong, L. (2009) Expression of Ebolavirus glycoprotein on the target cells enhances viral entry. Virol. J. 6, 75

37. Brandt, J., Wendt, L., and Hoenen, T. (2019) Structure and functions of the Ebola virus matrix protein VP40. Future virology 14, 21–30

38. Bank, P. D. (1971) Protein data bank. Nature New Biol. 233, 223

39. Eswar N., Webb, B., Marti-Renom, M. A., Madhusudhan, M. S., Eramian, D., Shen, M-y., et al. (2006) Comparative protein structure modeling using Modeller. Curr. Protoc. Bioinformatics 5, 5.6

40. Jo, S., Kim, T., Iyer, V. G., and Im, W. (2008) CHARMM-GUI: a web-based graphical user interface for CHARMM. J Comput Chem 29, 1859–1865

41. Wu, E. L., Cheng, X., Jo, S., Rui, H., Song, K. C., Davila-Contreras, E. M., et al. (2014) CHARMM-GUI Membrane Builder toward realistic biological membrane simulations. J Comput Chem 35, 1997–2004

42. Zachowski, A. (1993) Phospholipids in animal eukaryotic membranes: transverse asymmetry and movement. Biochem. J. 294 ( **Pt 1**), 1–14

43. van Meer, G., Voelker, D. R., and Feigenson, G. W. (2008) Membrane lipids: where they are and how they behave. Nat. Rev. Mol. Cell Biol. 9, 112–124

44. Brooks, B. R., Brooks, C. L., 3rd, Mackerell, A. D., Jr., Nilsson, L., Petrella, R. J., Roux, B., et al. (2009) CHARMM: the biomolecular simulation program. J Comput Chem 30, 1545–1614

45. Phillips, J. C., Hardy, D. J., Maia, J. D., Stone, J. E., Ribeiro, J. V., Bernardi, R. C. et al. (2020) Scalable molecular dynamics on CPU and GPU architectures with NAMD. J. Chem. Phys. 153, 044130

46. Mackerell, A. D., Jr. (2004) Empirical force fields for biological macromolecules: overview and issues. J. Comput. Chem. 25, 1584–1604

47. Essmann, U., Perera, L., Berkowitz, M. L., Darden, T., Lee, H., and Pedersen, L. G. (1995) A smooth particle mesh Ewald method. J. Chem. Phys. 103, 8577–8593

48. Ryckaert, J.-P., Ciccotti, G., and Berendsen, H. J. C. (1977) Numerical integration of the cartesian equations of motion of a system with constraints: molecular dynamics of n-alkanes. J. Comp. Phys. 23, 327–341

49. Brooks, M. M., Hallstrom, A., and Peckova, M. (1995) A simulation study used to design the sequential monitoring plan for a clinical trial. Stat. Med. 14, 2227–2237

50. Humphrey, W., Dalke, A., and Schulten, K. (1996) VMD: visual molecular dynamics. J. Mol. Graph. 14, 33–38

51. Del Vecchio, K., and Stahelin, R. V. (2016) Using surface plasmon resonance to quantitatively assess lipid–protein interactions. Lipid Signaling Protocols 1376, 141–153

52. Nanbo, A., Maruyama, J., Imai, M., Ujie, M., Fujioka, Y., Nishide, S. et al. (2018) Ebola virus requires a host scramblase for externalization of phosphatidylserine on the surface of viral particles. PLoS Pathog. 14, e1006848

53. Kuroda, M., Fujikura, D., Nanbo, A., Marzi, A., Noyori, O., Kajihara, M. et al. (2015) Interaction between TIM-1 and NPC1 is important for cellular entry of Ebola virus. J. Virol. 89, 6481–6493

