## Supplemental Figures for "Minor changes in electrostatics robustly increase VP40 membrane binding, assembly, and budding of Ebola virus matrix protein derived virus-like particles"

Short title: Electrostatic changes alter VP40 assembly and budding

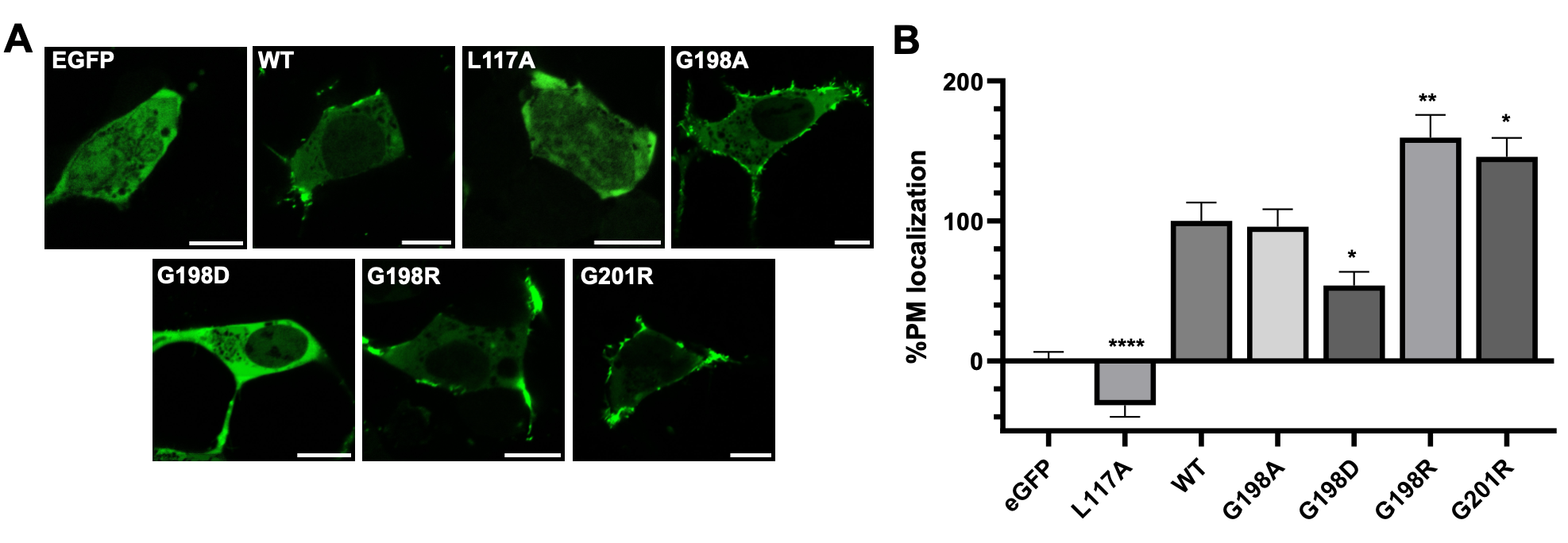

**Figure S1. Cellular localization after eight hours for WT VP40 and respective mutants in HEK293 cells.** *A*, Representative images of EGFP, EGFP-WT-VP40, EGFP-VP40-L117A, EGFP-VP40-G198A, EGFP-VP40-G198D, EGFP-VP40-G198R, EGFP-VP40-G201R constructs imaged 8 hours post-transfection. Scale bar = 10 μm. *B*, Average PM localization for each construct. Image analysis was done in Image J to determine percent PM localization (%PM localization) for each construct. %PM localization was defined as (PM intensity/(PM intensity+ cytosol intensity))*100. Data was normalized to %PM localization of WT as 100% localization and %PM localization of EGFP as 0% localization. A WGA-Alex647 PM marker was used to mark the PM in each cell. Error bars are the SEM from three independent experiments with at least 9 images analyzed per replicate. A one-way ANOVA was performed with multiple comparisons compared with WT PM localization ( *p<0.05, **p<0.01 and ****p<0.0001).

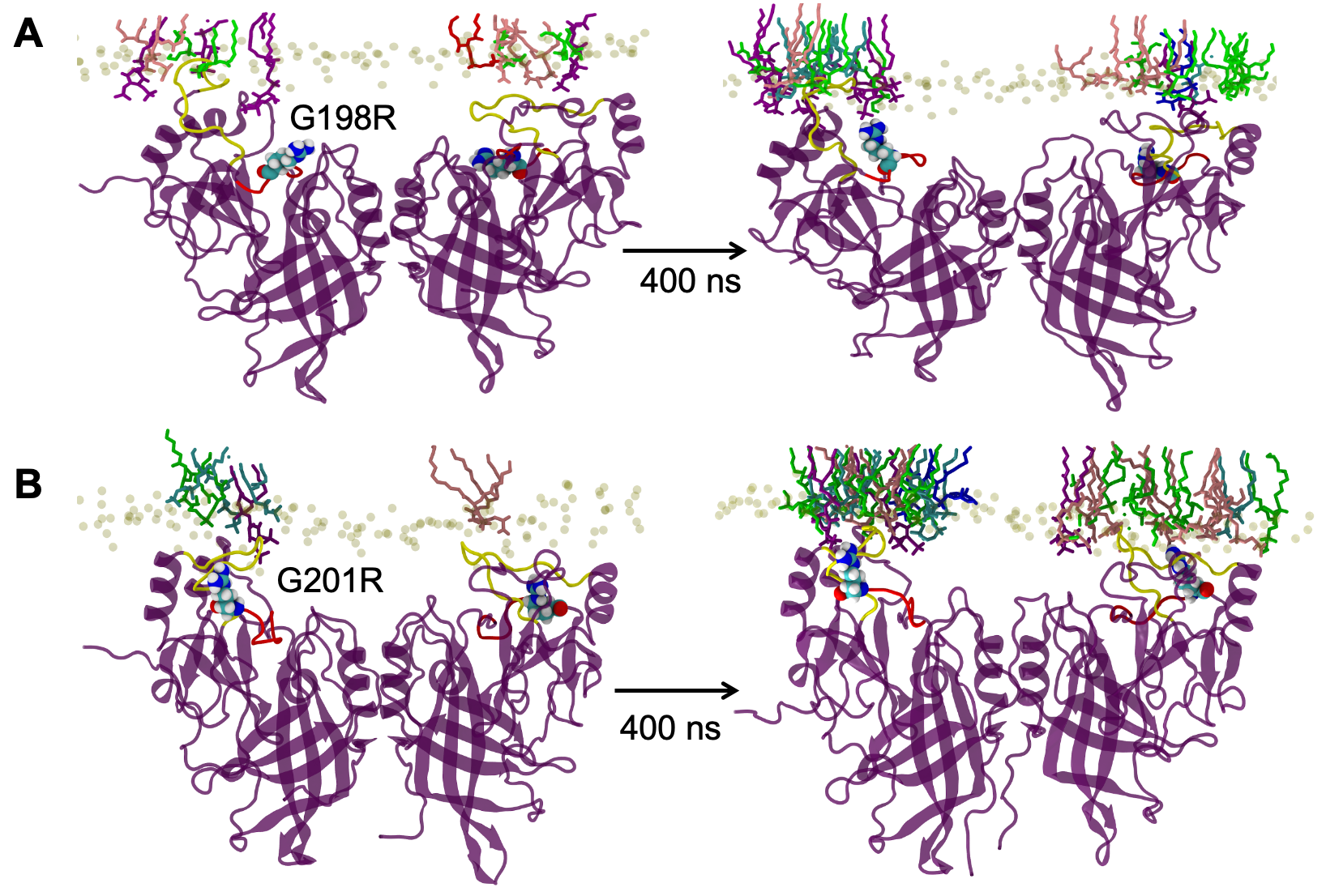

**Figure S2 . Snapshots of G198R and G201R molecular dynamics simulations reveal increased C-terminal domain electrostatic interactions with the membrane at 400 ns.** *A*, G198R increased membrane contacts between VP40 and the membrane interface compared to WT VP40 during the simulation. This includes increased contacts by R198, K221, K224, N227 and K279 residues. *B*, G201R leads to increased electrostatic association with the anionic membrane compared to WT as evidenced by increased interactions by residues K104, R201, K224, and S278.

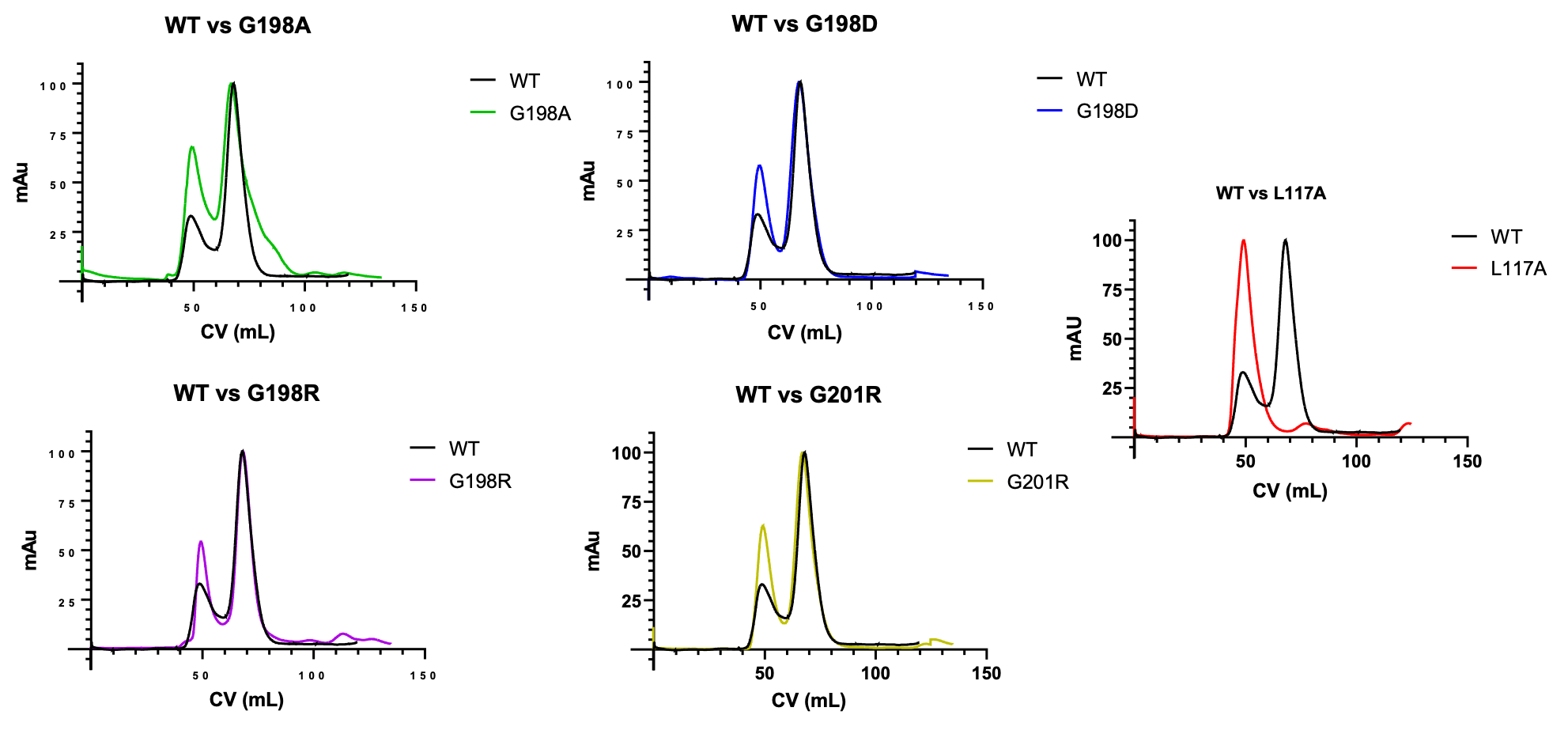

**Figure S3. Dimer and octamer assessment of WT VP40 and mutations using size exclusion chromatography.** Normalized size exclusion chromatogram trace of WT, G198A, G198D, G198R and G201R purified VP40 protein shows similar formation of the dimer in solution compared to WT (only dimers used for lipid-binding assays). L117A displayed an increased propensity towards the octamer form compared to WT VP40 and a smaller peak (at ~78 mL CV) we suspect is the monomeric form of VP40 (monomer used for lipid-binding assays). Please note the same WT VP40 purification profile is shown in each graph.

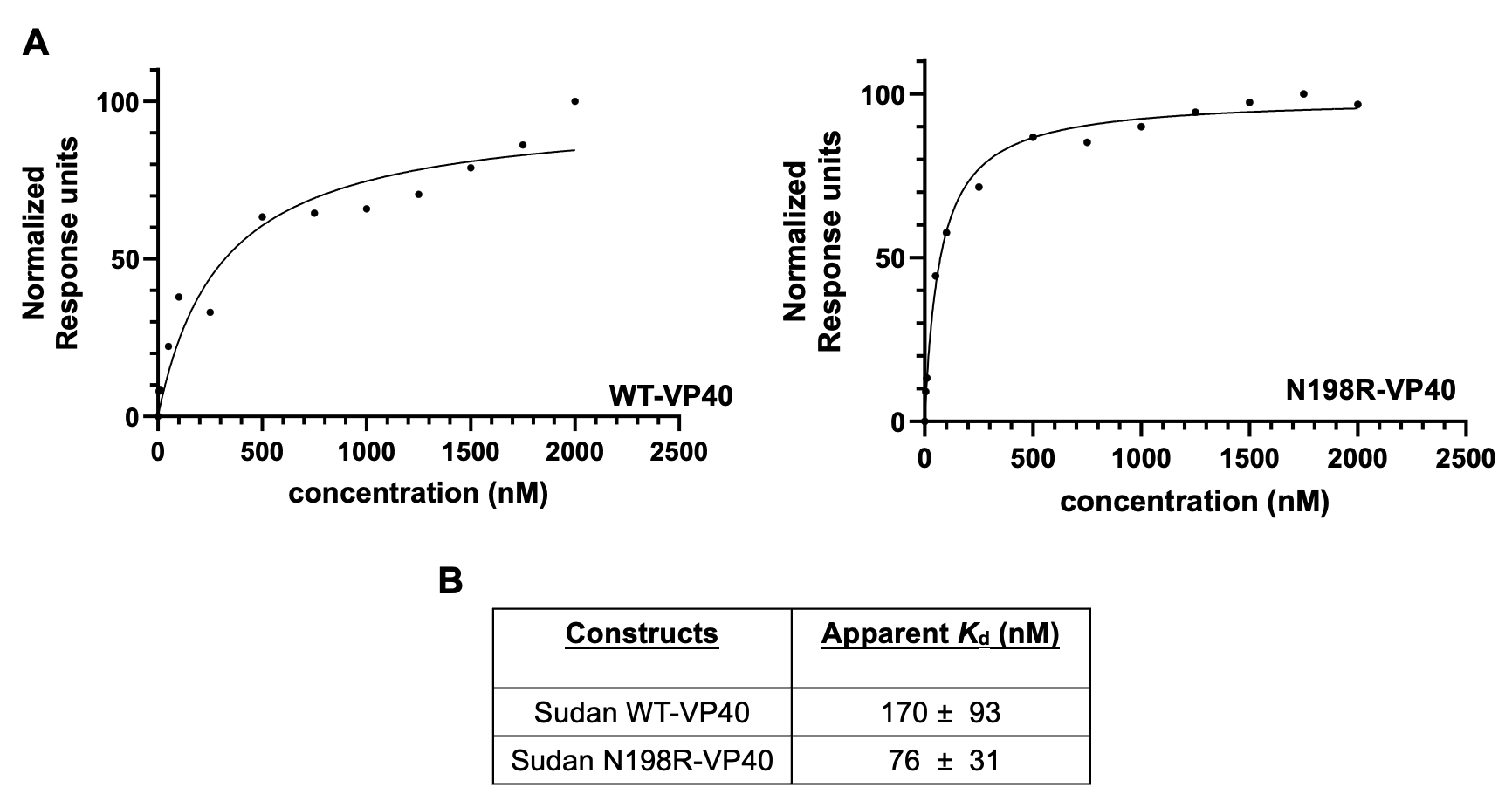

**Figure S4. SPR binding measurements for Sudan ebolavirus WT delta 43 VP40 and delta 43 N198R VP40 for lipid vesicles containing POPS**. *A,* Representative SPR binding curves for WT Sudan VP40 and N198R VP40 to LUVs (POPC:POPE:POPS (60:20:20)) containing 20% POPS, n = 4 for WT VP40 and n = 3 N198R VP40 measurements. *B*, Table showing the apparent binding affinities (*K* _d_) and STDEV of WT VP40 and mutant to lipid vesicles containing 20% POPS.

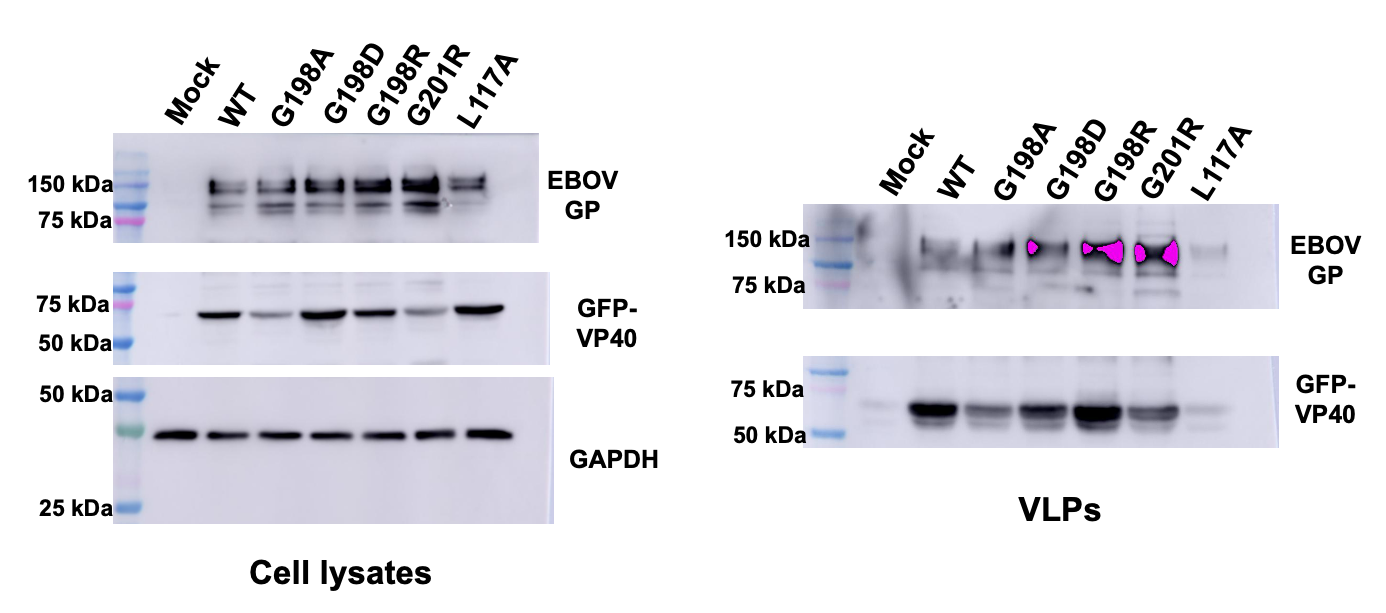

**Figure S5. Representative Western blot for cell lysates and VLPs collected from EBOV GP-VP40 co-expressed with WT VP40 or respective VP40 mutations expressed in HEK293 cells.** HEK293 cells were transfected with plasmid DNA encoding EGFP-WT-VP40, EGFP-VP40-L117A, EGFP-VP40-G198A, EGFP-VP40-G198D, EGFP-VP40-G198R, or EGFP-VP40-G201R. The VLPs and cell lysates were collected 48 hrs after transfection and subjected to Western blot using anti-GFP. GAPDH was used as a loading control for these experiments. Molecular weight markers are shown for all blots to verify band size for EGFP-VP40 and GAPDH.
